# Natural variation in the *irld* gene family affects starvation resistance in *C. elegans*

**DOI:** 10.1101/2021.06.07.447366

**Authors:** Amy K. Webster, Rojin Chitrakar, Maya Powell, Jingxian Chen, Kinsey Fisher, Robyn Tanny, Lewis Stevens, Kathryn Evans, Angela Wei, Igor Antoshechkin, Erik C. Andersen, L. Ryan Baugh

## Abstract

Starvation resistance is important to disease and fitness, but the genetic basis of its natural variation is unknown. We developed a synthetic-population (re)sequencing approach using molecular inversion probes (MIP-seq) to measure relative fitness during and after larval starvation in *C. elegans*. We applied this competitive assay to 100 genetically diverse, sequenced, wild strains, revealing natural variation in starvation resistance. We confirmed that the most starvation-resistant strains survive and recover from starvation better than the most starvation-sensitive strain, MY2147, using standard assays. We performed genome-wide association with the MIP-seq trait data and identified three quantitative trait loci (QTL) for starvation resistance. These QTL contain several members of the Insulin/EGF Receptor-L Domain (*irld*) family with sequence variation associated with variation in starvation resistance. We used genome editing to show that individual modification of four *irld* genes increases starvation resistance of MY2147. Modification of *irld-39* and *irld-52* together increases starvation resistance of the laboratory-reference strain N2. Increased starvation resistance of the *irld-39; irld-52* double mutant depends on *daf-16/FoxO*, and these worms also show increased nuclear localization of DAF-16 during starvation. DAF-16/FoxO is a widely conserved transcriptional effector of insulin/IGF signaling (IIS), and these results suggest that IRLD proteins modify IIS. This work demonstrates efficacy of using MIP-seq to dissect a complex trait, identifies *irld* genes as natural modifiers of starvation resistance in *C. elegans*, and suggests that an expanded gene family affects a deeply conserved signaling pathway to alter a fitness-proximal trait.

## INTRODUCTION

Enduring periods of starvation is a near-ubiquitous feature of animal life that affects survival, growth, and reproduction, making starvation resistance a fitness-proximal trait. Starvation resistance is also important to human health and disease, with direct relevance to diabetes, obesity, aging, and cancer. However, the genetic basis of natural variation in starvation resistance is unclear, despite its importance to understanding animal evolution and informing therapeutic strategies.

The nematode *C. elegans* is frequently starved in the wild and has robust starvation responses (Schulenburg and FÉlix 2017; Baugh and Hu 2020). Larvae that hatch in the absence of food arrest development in the first larval stage (L1 arrest or L1 diapause) and can survive starvation for weeks (Baugh 2013). In addition to causing mortality, extended starvation reduces growth and reproductive success upon feeding (Jobson *et al*. 2015; Jordan *et al*. 2019), and these effects can be uncoupled (Baugh and Hu 2020). For example, mutations affecting the endoplasmic reticulum unfolded-protein response do not affect starvation survival during L1 arrest, but they render starved worms incapable of growth after feeding (Roux *et al*. 2016). Likewise, sensory perception of food during L1 starvation does not affect survival but causes irreversible developmental arrest (Kaplan *et al*. 2018). In addition, *daf-18/PTEN* mutant worms display reduced L1 starvation survival and growth upon recovery, but their reproduction is particularly sensitive to starvation, with only 12 hours of L1 starvation causing dramatic decline in brood size, including sterility in some individuals (Chen *et al*. 2022). Starvation resistance therefore integrates survival, growth rate, and reproductive success, and different genes and conditions can affect these three phenotypes independently.

Insulin signaling is reduced during starvation, and evolutionary biologists, geneticists, and clinicians have all noted the parallel roles of insulin signaling in metabolic syndrome and starvation resistance (Soeters and Soeters 2012; Flier 2019). Insulin/IGF signaling (IIS) is a critical regulator of *C. elegans* L1 arrest (Baugh and Sternberg 2006). There is a single known insulin/IGF-like receptor in *C. elegans*, DAF-2/InsR, which signals through a conserved phosphatidylinositol 3-kinase (PI3K) signaling pathway to antagonize the forkhead boxO transcription factor DAF-16/FoxO (Lin *et al*. 1997; Ogg *et al*. 1997). When IIS is active, DAF-16 is phosphorylated by AKT-1/AKT, sequestering it in the cytoplasm, and when IIS is reduced, such as during starvation, DAF-16 moves to the nucleus and regulates transcription (Henderson and Johnson 2001; Lee *et al*. 2001; Lin *et al*. 2001). *daf-16/FoxO* promotes starvation resistance, with *daf-16* and *daf-2* mutants displaying starvation sensitivity and resistance, respectively (Munoz and Riddle 2003; Baugh and Sternberg 2006; Hibshman *et al*. 2017)

IIS is subject to extensive regulation, presumably reflecting its critical role in maintaining metabolic and developmental homeostasis during fluctuations in nutrient availability. There are 40 insulin-like peptides (ILPs) encoded by the *C. elegans* genome, and they function as either agonists or antagonists of DAF-2/InsR (Pierce *et al*. 2001). There is evidence of functional redundancy, but genetic analysis also suggests functional specificity among ILPs beyond their broad classification as either agonists or antagonists (Fernandes de Abreu *et al*. 2014). ILPs are expressed in sensory and other neurons, the intestine, and other cell types (Ritter *et al*. 2013), and they contribute to systemic regulation (Chen and Baugh 2014; Hung *et al*. 2014). IIS and *daf-16/FoxO* regulate transcription of most ILPs, producing extensive feedback regulation (Kaplan *et al*. 2019), and comprising a putative insulin-to-insulin signaling network (Fernandes de Abreu *et al*. 2014). The *daf-2/InsR* locus encodes a truncated transcript isoform, *daf-2b*, that produces a product lacking a cytoplasmic domain such that it is incapable of signaling. Instead, it presumably binds ILPs, competing with full-length DAF-2 isoforms (Martinez *et al*. 2020). In a search for other potential insulin/IGF receptors, *C. elegans* proteins with homology to various insulin receptors were identified by PSI-BLAST, leading to the discovery of a large family of proteins with weak homology to DAF-2/InsR (Dlakic 2002). These extracellular proteins lack a tyrosine kinase domain, like DAF-2B, but they have an atypical ligand-binding domain that also shares homology with epidermal growth factor (EGF) receptors. 68 members of this Insulin/EGF-<u>R</u>eceptor <u>L D</u>omain (IRLD) protein family are now recognized in *C. elegans* (Hobert 2013). It was originally hypothesized that the IRLD proteins function as decoy insulin/IGF receptors (Dlakic 2002), similar to how DAF-2B is thought to function (Martinez *et al*. 2020), but genetic analysis of two of them (*hpa-1* and *hpa-2*) suggests that they modify EGF signaling to affect healthspan (Iwasa *et al*. 2010). The IRLD family is not conserved outside of the genus (Oliver Hobert, personal communication), but there is a family of IGF-binding proteins in vertebrates that is thought to fine-tune IGF signaling by affecting circulation and receptor binding of IGF proteins (Allard and Duan 2018).

Hundreds of genome-sequenced, wild *C. elegans* strains are available for genetic analysis of natural variation on the *Caenorhabditis elegans* Natural Diversity Resource (CeNDR) (Cook *et al*. 2017). Short-read sequencing was mapped to the N2 reference genome, and millions of single-nucleotide variants (SNVs) were called. Analysis of these SNVs in conjunction with read mapping efficiency for each strain led to the identification of hyper-divergent regions of the genome, which exhibit substantially more variation than other regions of the genome for some strains (Lee *et al*. 2021). These regions are typically found on the arms of chromosomes and are enriched for environmental-response genes.

We were interested in taking advantage of the genomic resources on CeNDR to investigate natural variation of starvation resistance. Assaying starvation resistance for so many strains is prohibitively laborious using traditional methods. We previously developed an approach based on pooling wild strains, selecting for starvation resistance, and sequencing to identify relatively starvation-resistant and sensitive strains (Webster *et al*. 2019). This approach identified a quantitative trait locus (QTL), but it lacked power since only very rare sequencing reads that include SNVs unique to a strain in the synthetic population informed inference of relative strain frequency. In contrast, by capturing targeted sequences, <u>m</u>olecular inversion <u>p</u>robe sequencing (MIP-seq) enables extremely deep sequencing of polymorphic loci (Cantsilieris *et al*. 2017; Mok *et al*. 2017), providing sensitive and precise estimation of strain frequencies in a complex pool as a means of high-throughput phenotyping. Here, we used MIP-seq to phenotype starvation resistance in a pool of genetically diverse wild *C. elegans* strains. We identified three QTL and four members of the *irld* gene family that affect starvation resistance, and our results suggest that they do so at least in part by modifying IIS.

## RESULTS

### Sensitive and precise measurement of strain frequency in pooled culture using MIP-seq

We selected 103 genetically diverse, wild *C. elegans* strains from around the world including the laboratory reference N2 to test MIP-seq, ultimately phenotyping 100 for starvation resistance (Fig 1A-B). Molecular inversion probes (MIPs) are designed to capture a specific region of the genome for targeted multiplex sequencing (Fig 1C). We designed MIPs to target a region containing a SNV unique to each of 103 strains. Thus, the relative frequency of each strain in a pool can be determined by the SNV frequency. We designed four such MIPs per strain to provide redundancy and increase precision. To pilot MIP-seq, we prepared sequencing libraries from an equimolar mix of genomic DNA from each of 103 strains. We determined the frequency of strain-specific reads for each MIP, and we censored probes that produced frequencies substantially different than the expected value of approximately 0.01 (Fig 1D; criteria in Materials and Methods), leaving three or four reliable probes for 85% of strains and at least one MIP for 100 strains (Fig 1E, Data S1), which were used in the starvation resistance experiment. Three strains with no MIPs passing filtering were excluded from subsequent analysis. As an additional pilot, we mixed genomic DNA from a subset of strains at different concentrations to prepare a standard curve. MIP-seq accurately measured individual strain frequencies over three orders of magnitude (Fig 1F), and greater sequencing depth could theoretically expand the dynamic range.

**Fig. 1.**
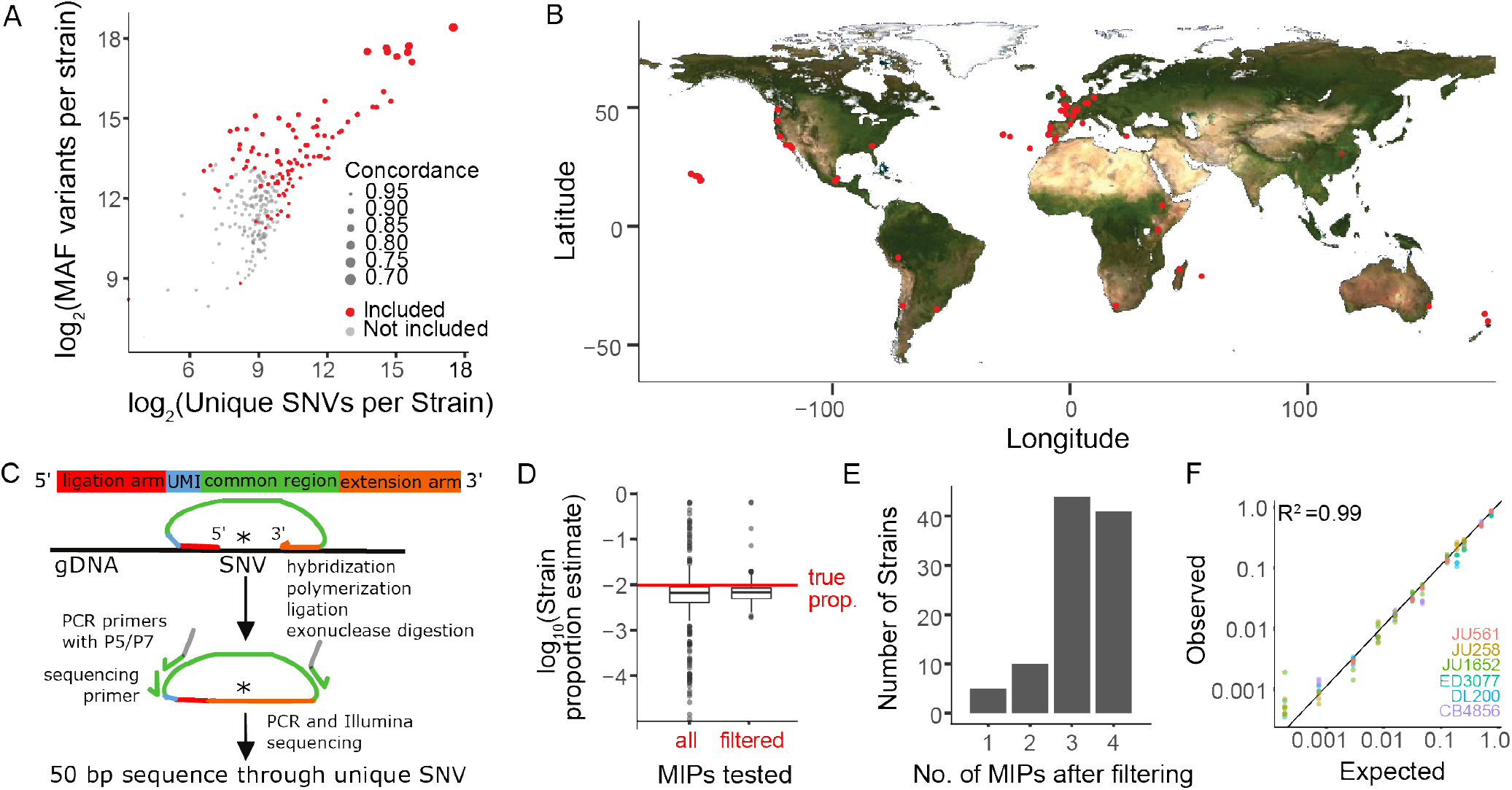
Sensitive and precise measurement of strain frequency in pooled culture using MIP-seq. (**A**) The three metrics used to identify the most diverse *C. elegans* strains are plotted. “MAF” stands for minor allele frequency. Concordance refers to the average pairwise concordance for the focal strain compared to all other strains, which is calculated as the number of shared variant sites divided by the total number of variants for each pair. The strains included in the MIP-seq experiments are in red. (**B**) Geographic locations of the strains assayed for starvation resistance. (**C**) Schematic of MIP-seq. MIPs are designed for loci with SNVs unique to each strain. Four MIPs were designed per strain. MIPs are 80 nt long and include ligation and extension arms to match DNA sequence surrounding the SNV, a unique molecular identifier (UMI), and P5 and P7 sequences for Illumina sequencing. MIPs are hybridized to genomic DNA, polymerized, ligated, and used as PCR template to generate an Illumina sequencing library. The alternative-to-total read frequency for each MIP/SNV locus indicates strain frequency. (**D**) Empirical testing of 412 MIPs with an equimolar mix of genomic DNA from 103 strains to identify reliable MIPs. 321 MIPs passed filtering and were analyzed in the starvation experiment. Outliers for filtered MIPs are for N2, which has hardly any unique SNVs because it is derived from the reference genome. N2 MIPs were included despite poor performance. (**E**) Number of MIPs per strain of the 321 filtered MIPs that passed filtering. (**F**) Genomic DNA from 7 strains was combined at different known concentrations, and MIP-seq was used to generate a standard curve. Included MIPs all passed filtering. (R^2^ = 0.99).

### Using MIP-seq to characterize natural variation of starvation resistance

We used MIP-seq to phenotype 100 diverse strains for starvation resistance. We cultured the strains in standard laboratory conditions, pooled them, and subjected them to starvation during L1 arrest. We aimed for approximately 5,000 L1 larvae per strain in the pooled starvation culture in order to ensure representative sampling. However, we expected actual representation to vary to some extent across strains and replicates, so we collected DNA from an aliquot of L1 larvae on the first day of starvation as a “baseline” sample to capture initial population composition. In addition, aliquots were taken from the pool at days 1, 9, 13, and 17 of starvation, and sampled larvae were allowed to recover with food for four or five days (depending on the duration of starvation), enabling reproduction for one day, and then the entire population was collected for DNA preparation (Fig 2A). DNA from baseline samples, as well as samples allowed to recover and reproduce following starvation, were sequenced with MIP-seq for each of five biological replicates. It is critical to point out that by incorporating recovery and early fecundity, this sampling scheme integrates effects of starvation on mortality as well as growth rate and reproductive success, each of which are important for starvation resistance (*i.e*., fitness) and can be uncoupled by certain genotypes and conditions.

**Fig. 2.**
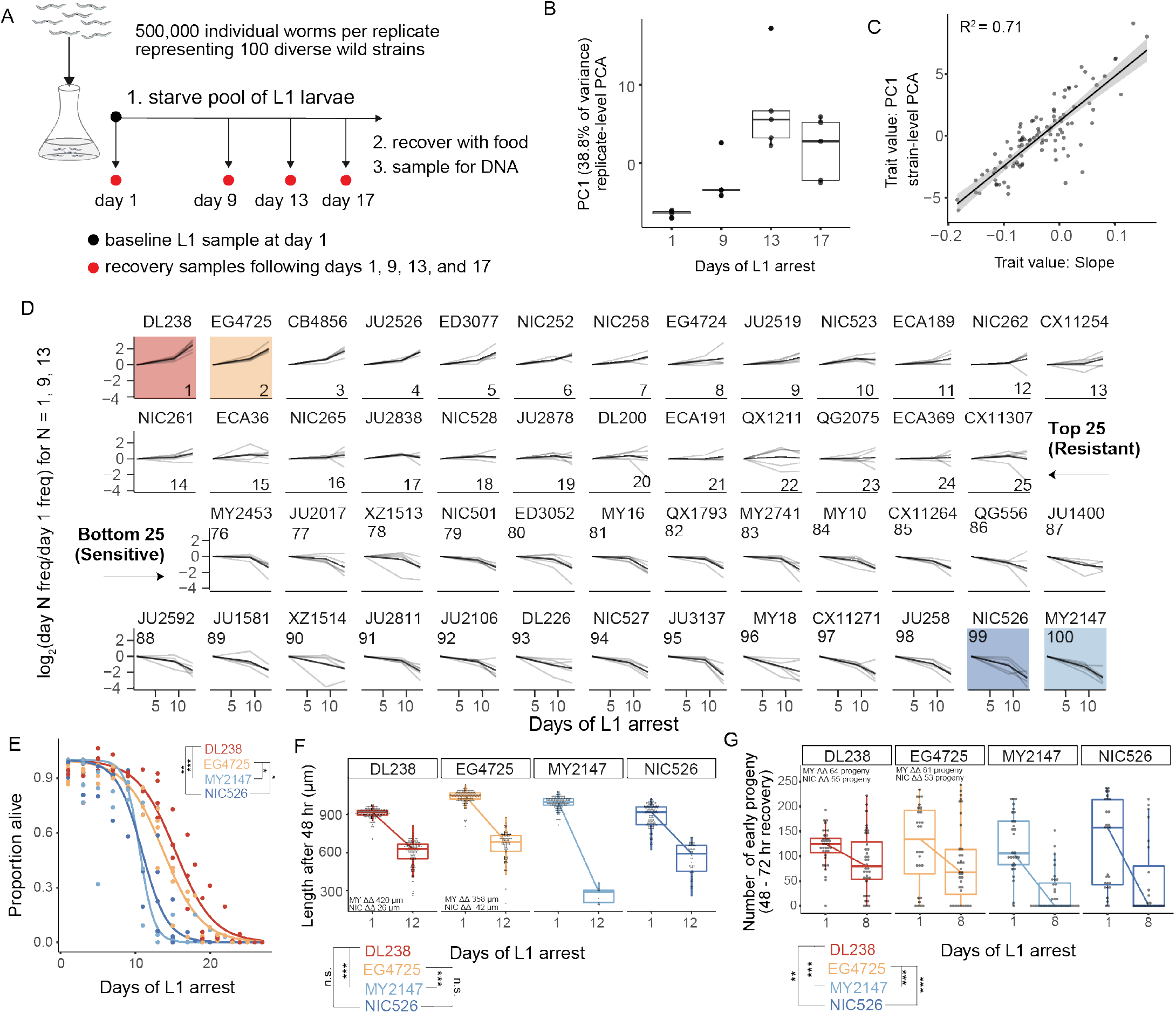
MIP-seq determines relative starvation resistance of 100 strains. (**A**) Experimental design. Worms were starved at the L1 stage (’L1 arrest’). ∼5,000 L1 larvae per strain were starved (∼500,000 total). The population of starved L1 larvae was sampled initially (”baseline” on day 1), and then sampled on the days indicated. Samples (except baseline) were recovered with food in liquid culture, reaching adulthood and producing progeny for one day, and the entire population was frozen for DNA isolation. Five biological replicates were performed. (**B**) Principal component 1 of normalized and processed data from all replicates (replicate-level) and strains is plotted, revealing association with duration of starvation. Each point is an individual sample (MIP-seq library). (**C**) The relationship between two starvation-resistance metrics (Slope and PC1) produced from strain-level data (replicates averaged) is plotted. Each point is a different strain. (**D**) Log_2_-normalized strain frequency is plotted over time for the 25 most resistant and 25 most sensitive strains in rank order (based on Slope). Only days 1, 9, and 13 are plotted. See Supplementary Figure 2 for full data. Grey lines are biological replicates and black line is the mean. DL238 and EG4725 are most starvation-resistant, and NIC526 and MY2147 are most sensitive, and they are color-coded accordingly. (**E**) L1 starvation survival curves are plotted for starvation-resistant and sensitive strains. Individual replicate measurements are included as points to which curves were fit with logistic regression. T-tests on 50% survival time of four biological replicates. (**F**) Worm length following 48 hours of recovery with food after 1 or 12 days of L1 starvation. (**G**) Number of progeny produced between 48 and 72 hours of recovery on food following 1 or 8 days of starvation. (**F,G**) ΔΔ indicates effect size of interaction between duration of starvation and strain data plotted in that panel compared to the strain listed (the difference in differences between strains’ mean length at days 1 and 12 or between mean number of progeny at days 1 and 8). ‘MY’ is an abbreviation for MY2147 and ‘NIC’ is an abbreviation for NIC526. Linear mixed-effects model; one-way p-value of interaction between duration of starvation and strain. (**E-G**) ***p < 0.001, **p < 0.01, *p < 0.05

DNA from ‘baseline’ samples allowed us to effectively normalize differences in pool composition in each replicate, revealing effects of starvation on strain frequency. Differences in pool composition explained the first principal component in principal component analysis (PCA) when strain frequencies over time were analyzed without consideration of baseline frequencies (Fig S1A). However, once the data were normalized for initial strain composition using the baseline sample for each replicate, the first principal component correlated with duration of starvation, especially across the first three time points (Fig 2B, Fig S1B). Substantial mortality occurred by day 17 (Fig S1C), and day 17 recovery cultures thus produced relatively few progeny. Consequently, differences in strain frequencies were actually smaller at day 17 than 13, but relative differences were conserved (Fig S2). Therefore, after normalization, duration of starvation is the major factor accounting for differences in strain frequency across all samples, and this is robust to differences in the initial composition of the pool across replicates.

We developed two metrics to quantify relative starvation resistance for each strain. “Slope” is a measure of how much a strain increases or decreases in frequency over time across days 1, 9, and 13, calculated as the slope of a linear model (Data S1). “PC1” is the value of the first principal component for each strain from strain-level PCA (Fig 2B, Fig S1B). These two metrics are correlated but also show some differences (Fig 2C), suggesting they capture related but also distinct features of the data. While Slope is intuitive, it is limited by the use of a linear model. Nonetheless, Slope values are correlated with starvation-resistance values produced from a previously published population-sequencing approach with less power that included some of the same strains (Fig S1D) (Webster *et al*. 2019). In addition, Slope is modestly correlated with the latitude from which strains were collected, suggesting possible adaptation to starvation or other correlated traits based on location (Fig S3). There is also a modest negative correlation between Slope and growth rate after only one day of starvation (control condition) (Fig S4), suggesting a possible trade-off between starvation resistance and population growth rate in the absence of stress. We used the Slope metric to order strains from most resistant to sensitive, revealing differences in starvation resistance between wild strains (Fig 2D, Fig S2). In contrast to Slope, PC1 does not assume linearity, it includes the results from day 17 of starvation, and it may be less affected by noise. PCA is also a common way to obtain trait values for genome-wide association (GWA) studies (Ried *et al*. 2016; Yano *et al*. 2019).

Our recovery-based sampling approach integrates starvation survival, recovery, and early fecundity into a single fitness assay. It is therefore unclear whether a given strain is more or less resistant because of differences in mortality, growth rate, or progeny production. For example, a particular strain could survive no better than others, but it could grow faster upon recovery or retain greater reproductive success in order to display relative resistance to starvation. Conversely, a strain may not survive as long as others despite comparable growth or fecundity upon recovery in order to display relative sensitivity to starvation. It is also unclear what the absolute effect sizes are between the most resistant and sensitive strains in this competition assay. Nonetheless, our approach is intended to model the impact of larval starvation on fitness broadly, while traditional assays can be used to isolate specific effects of starvation on survival, growth, and reproduction in follow-up experiments.

We performed manual assays for starvation survival, growth rate, and early fecundity for the most resistant and sensitive strains. We found starvation-resistant strains DL238 and EG4725 survived starvation significantly longer during L1 arrest than sensitive strains MY2147 and NIC526 (Fig 2E). Differences in starvation survival among wild strains are relatively small compared to many published mutants in the N2 reference background (Baugh and Hu 2020), suggesting that large-effect mutations are purged by selection. After extended L1 arrest, DL238 and EG4725 recovered from starvation better than MY2147 but not NIC526, as assessed by their size following 48 hours of recovery (Fig 2F). Finally, DL238 and EG4725 exhibited a larger early brood size following extended starvation compared to both MY2147 and NIC526. Overall, this demonstrates that differences in starvation resistance among wild strains are driven by differences in survival, recovery, and early fecundity, but that sensitivity of NIC526 is apparently driven by differences in survival and early fecundity without an appreciable effect on growth. These results validate the MIP-seq approach and reveal the extent of natural variation in starvation resistance.

### Natural variation in *irld* gene family members affects starvation resistance

We used Slope and PC1 as trait values to perform GWA on CeNDR (Cook *et al*. 2017). GWA identified QTL on the right arm of chromosome IV and on the left and right arms of chromosome V (Fig 3A-B, Data S2). We confirmed that each QTL affected starvation resistance by generating near-isogenic lines and measuring growth rate upon recovery from starvation (Fig S5). We chose this assay, as opposed to starvation survival or early fecundity, because it revealed relatively robust differences between DL238/EG4725 (resistant) and MY2147 (sensitive) (Fig 2F).

**Fig. 3.**
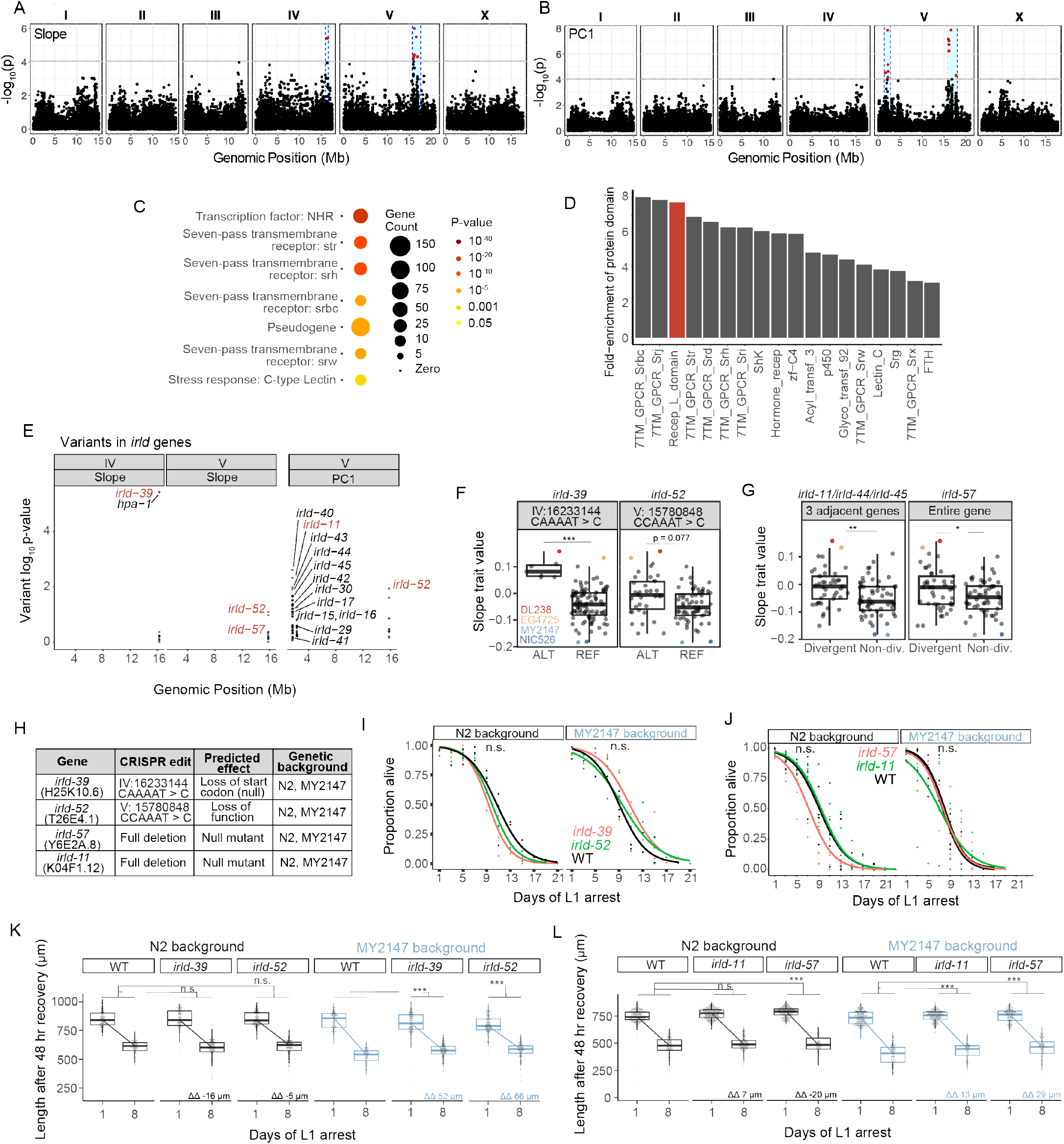
Genetic variation in the *irld* gene family underlies differences in starvation resistance. (**A**) GWA output using Slope as a trait value. Significant QTL intervals are IV: 15939340-16613710 and V: 15660911-17615557. (**B**) GWA output using PC1 as a trait value. Significant QTL intervals are V: 1345848-2764788 and V: 15775895-18065050. (**C**) WormCat Category 3 enrichments for all genes with variants in the QTL. (**D**) Fold-enrichment of protein domains significantly enriched among genes with variants in QTL. A hypergeometric p-value was calculated for each of 102 protein domains present, and a Bonferroni-corrected p-value of 0.00049 was used as a threshold to determine significance. Red indicates the receptor L domain, which is found in *irld* genes. (**E**) All variants in *irld* genes that are within significant QTL and their association with the starvation-resistance traits, Slope and PC. Each gene name is shown next to the most significant variant for that gene, but multiple variants are plotted for each gene when present. Red indicates genes selected for functional validation. (**F**) Slope trait values for strains based on whether they have ALT and REF alleles for specific *irld-39* and *irld-52* variants predicated to disrupt protein function. The *irld-52* variant p-value is p = 0.007 for the PC1 trait value (only the Slope trait value is shown here). Significance determined from GWA fine mapping. (**G**) Slope trait values for strains based on whether they are hyper-divergent or not at *irld-11* and *irld-57* loci. T-test on trait values between hyper-divergent and non-divergent strains. (**F-G**) DL238, EG4725, NIC526, and MY2147 are color-coded as indicated. (**H**) The four *irld* genes selected for genome editing and the edits generated for each. For *irld-39 and irld-52,* N2 and MY2147 have the REF allele and were edited to have the ALT allele. *irld-11* and *irld-57* are hyper-divergent in DL238 and EG2745 backgrounds, so full gene deletions were generated in N2 and MY2147 backgrounds. (**I**) L1 starvation survival assays on *irld-39* and *irld-52* ALT alleles in N2 and MY2147 backgrounds. There were no significant differences between strains within a background. (**J**) L1 starvation survival assays on *irld-11* and *irld-57* deletions in N2 and MY2147 backgrounds. There were no significant differences between strains within a background. (**K-L**). Worm length following 48 hr recovery with food after 1 or 8 d of L1 starvation for indicated genotypes. Linear mixed-effects model; one-way p-value for interaction between strain and duration of starvation; 4-5 biological replicates per condition. ΔΔ indicates effect size of interaction between duration of starvation and strain compared to control (the difference in differences between strains’ mean length at days 1 and 8). (**F,G,K,L**) ***p < 0.001, **p < 0.01, *p < 0.05, n.s. not significant.

These QTL include enrichments for several large gene families. WormCat analysis identified significant enrichments of serpentine receptors, nuclear hormone receptors, and C-type lectins (Fig 3C) (Holdorf *et al*. 2020). Likewise, protein-domain enrichment analysis (Finn *et al*. 2011) identified seven-pass transmembrane domains and hormone receptor domains (Fig 3D). In addition, the receptor L domain was significantly enriched, which is found in proteins comprising Insulin/EGF-Receptor L Domain (IRLD) family (Dlakic 2002). Given weak homology to DAF-2/InsR, and the critical role of IIS in regulation of starvation resistance, we were intrigued at the possibility that natural variation in *irld* family genes may impact starvation resistance. Across all three QTL identified, there are genetic variants in 16 *irld* genes, and 68 genes have been identified as part of this family in *C. elegans* (Hobert 2013). Multiple variants are present for each *irld* gene, and variants differed in the degree to which they were associated with variation in starvation resistance (Fig 3E; each gene name is indicated for its most significantly associated variant).

We selected at least one *irld* gene from each QTL for functional analysis. We used multiple criteria to select *irld* genes, including each variant’s association with starvation resistance, the predicted functional consequence of each variant (favoring variants that clearly disrupt protein function; for example, by introducing an early stop codon, frameshift, etc.), and its context among other variants. On the left arm of chromosome IV, a variant in *irld-39* was the strongest individual candidate both because of strong association with starvation resistance and because the variant is predicted to disrupt the start codon of the gene (Fig 3E, F, Data S2), likely rendering *irld-39* a functional null in the starvation-resistant strain DL238. On the right arm of chromosome V, *irld-52* was identified through both Slope and PC1 phenotype metrics and contains a variant associated with starvation resistance predicted to disrupt its fifth exon (Fig 3E, F). In this case, it is likely that protein function is disrupted but unclear if the variant causes a null mutation. We were thus interested in testing the function of both specific variants for *irld-39* and *irld-52* (Fig 3F). While analyzing variants on the left arm of chromosome V, we noticed that many *irld* genes are adjacent to each other and that each contain many variants, precluding identification of a single best candidate variant. In particular, *irld-11, irld-44,* and *irld-45* are directly adjacent to each other, and each gene contains over 50 genetic variants. This pattern of some loci containing many variants relative to N2 has been observed on a broader scale, leading to identification of so-called “hyper-divergent” regions of the genome that contain exceptional amounts of variation (Lee *et al*. 2021). A genomic region for which at least some strains are classified as hyper-divergent is considered a hyper-divergent region, but a given strain may or may not be hyper-divergent in a hyper-divergent region depending on its genomic sequence. *irld-11, irld-44,* and *irld-45* are part of a hyper-divergent region, and because they are in such close proximity, they are hyper-divergent in the same strains. We considered hyper-divergence status across these three genes as a summary of the variation in them, and found that hyper-divergence at these loci was associated with starvation resistance (Fig 3G). Finally, *irld-57* provides another example of a gene overlapping with a hyper-divergent region on the right arm of chromosome V, and hyper-divergence at this locus is also associated with starvation resistance (Fig 3G). We thus aimed to follow-up on the *irld* gene family by testing the functional consequences of variation in *irld-39, irld-52, irld-11,* and *irld-57.* Notably, the associations between variants or hyper-divergence and Slope (Fig 3F, G) together with their inferred negative impacts on gene function (Fig 3H) suggests that disruption of these *irld* genes in backgrounds where they are functional will increase starvation resistance.

We used CRISPR-Cas9 genome editing to determine functional consequences of genetic modification of our candidate *irld* genes. Because *irld-39* and *irld-52* contain singular variants associated with starvation resistance and predicted to disrupt protein function, we generated these specific variants in the starvation-sensitive MY2147 and the laboratory-reference N2 backgrounds (Fig 3H). Since *irld-11* and *irld-57* contain so many candidate variants, we deleted these genes in MY2147 and N2, rendering them null mutants at each locus (Fig 3H). We note that although the strains we generated are nulls for *irld-11* and *irld-57*, it is possible that the starvation-resistant wild strains may retain some function in these genes. In this sense, edits of *irld-39* and *irld-52* are more likely to approximate the effect of specific variants in the wild, because they are the exact variants present in starvation-resistant wild strains. We tested all eight strains and compared each to the appropriate control parental background, N2 or MY2147, to determine if these alleles affect starvation resistance. None of the alleles in either background significantly affected survival (Fig 3I, J). A power analysis suggests there is sufficient statistical power to detect differences of approximately two days or greater, suggesting there is not a difference of at least this magnitude. However, alleles for all four *irld* genes mitigated the effect of starvation on growth rate in the MY2147 background but not N2 (Fig 3K, L). This suggests that MY2147, as a more starvation-sensitive background than N2, facilitates detection of alleles that increase starvation resistance. In addition, these results suggest that these *irld* genes may uncouple survival and recovery from starvation to impact starvation resistance. It is also possible that we were able to detect an effect on starvation recovery simply because the assay is more sensitive. In summary, these results show that multiple types of variants in different *irld* family members reduce the effect of extended L1 starvation on recovery, identifying four individual genes from one family that affect starvation resistance in wild strains.

### IRLD-39 and IRLD-52 act through DAF-16/FoxO to affect starvation resistance

Because *irld-39(duk1)* and *irld-52(duk17)* alleles correspond to the exact variants present in starvation-resistant wild strains, we chose to focus on them for further analysis. We hypothesized that phenotypic effects may have been detected only in MY2147 because it is relatively sensitive to starvation, and multiple *irld* variants may reveal an effect in N2. An *irld-39(duk1); irld-52(duk17)* double mutant in the N2 background increased starvation survival but without statistical significance (Fig 4A). In this case there was sufficient statistical power to detect differences of approximately 1.5 days or greater, suggesting there is not a difference of at least this magnitude. However, the double mutant displayed a modest but significant increase in growth following 8 days of starvation (Fig 4B). Furthermore, the double mutant significantly increased early fecundity following starvation (Fig 4C). These results further support the conclusion that natural variation in *irld-39* and *irld-52* affects starvation resistance. Notably, the two variants are both present in the most starvation-resistant strain identified, DL238 (Fig 3F).

**Fig. 4.**
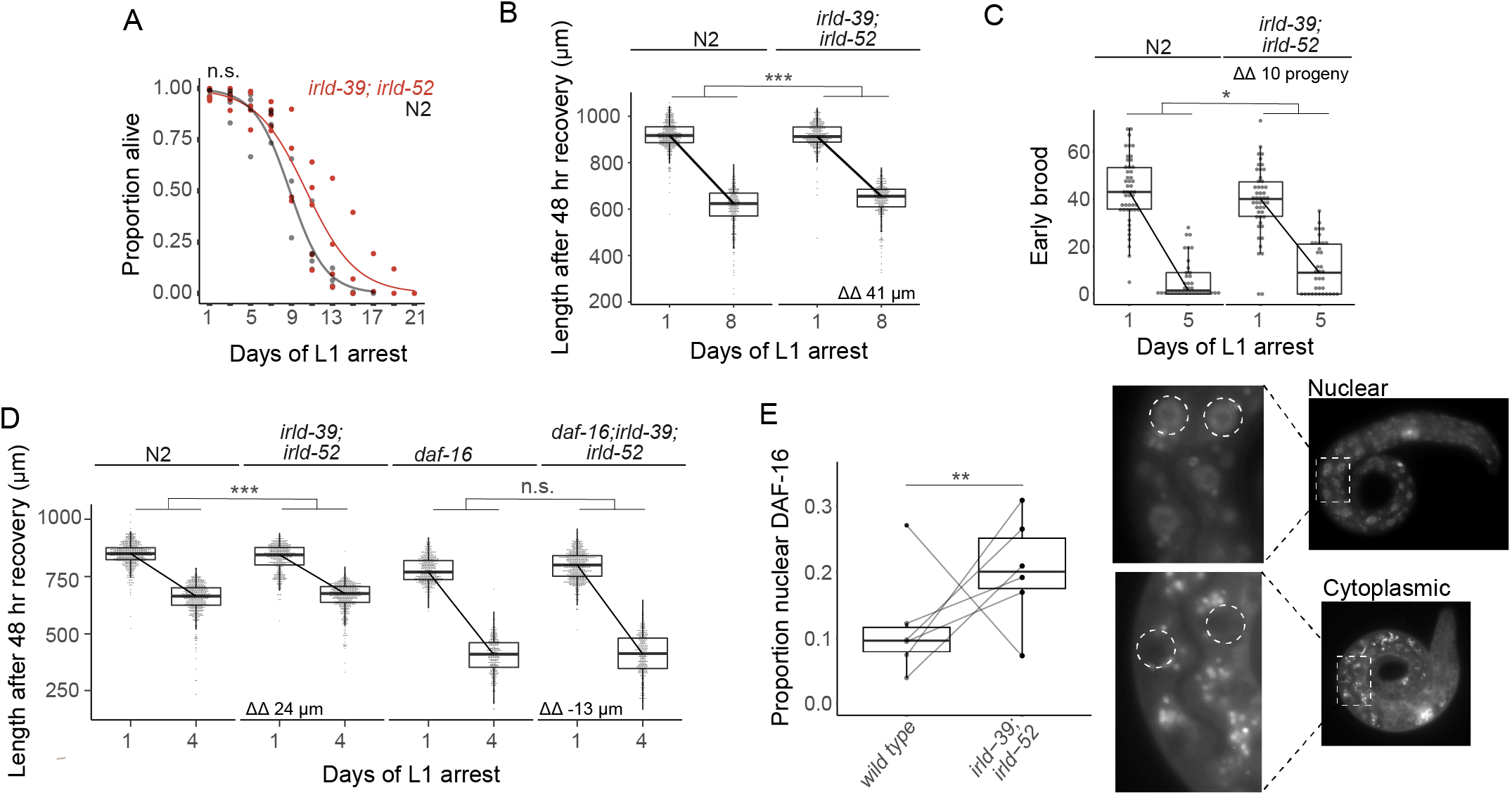
IRLD-39 and IRLD-52 together impact starvation resistance and depend on DAF-16. (**A**) Survival curves of *irld-39(duk1); irld-52(duk17)* and N2 throughout L1 starvation. The apparent increase in starvation survival in the double mutant is not statistically significant (p = 0.14). (**B**) Worm length of *irld-39(duk1); irld-52(duk17)* and N2 following 48 hours of recovery with food after 1 or 8 days of L1 starvation. (**C**) Number of progeny produced between 48 and 72 hours of recovery with food after 1 or 5 days of L1 starvation. (**D**) Worm length of N2, *irld-39(duk1); irld-52(duk17), daf-16(mu86),* and *daf-16(mu86); irld-39(duk1); irld-52(duk17)* following 48 hours of recovery with food after 1 or 4 days of L1 starvation. (**B-D**) Linear mixed-effects model with duration of L1 starvation and genotype as fixed effects and the number of replicates as a random effect; p-value calculated for interaction between fixed effects. ΔΔ indicates effect size of interaction between duration of starvation and strain compared to control. (**E**) Nuclear localization of DAF-16::GFP in intestinal cells of starved L1s ∼36 hours after hatching. Each point represents the result of a single independent biological replicate with 51-64 worms scored for each condition and replicate, with a line connecting the two genotypes in each replicate. The Cochran-Mantel-Haenszel test was used to determine differences in the distribution of the two categories (nuclear and cytoplasmic) between *daf-16(ot971)* (wild type) and *daf-16(ot971); irld-39(duk1); irld-52(duk17)* (*irld-39; irld-52*). Images of intestinal nuclear and cytoplasmic localization are shown. (**A-E**) 4-6 biological replicates were performed per experiment. ***p < 0.001, **p < 0.01, *p < 0.05, n.s. not significant.

Given weak homology between IRLD proteins and the extracellular domain of DAF-2/InsR, we wondered if IRLD-39 and IRLD-52 modify IIS, as originally proposed (Dlakic 2002). DAF-2/InsR activity inhibits the transcription factor DAF-16/FoxO, and *daf-2* mutants exhibit increased starvation resistance that is dependent on *daf-16* (Baugh and Sternberg 2006). We therefore hypothesized that increased starvation resistance with disruption of *irld-39* and *irld-52* depends on *daf-16/FoxO.* To test this, we generated a triple mutant including the null reference allele for *daf-16* (*daf-16(mu86); irld-39(duk1); irld-52(duk17)*). Notably, the *irld-39; irld-52* double mutant displayed significant mitigation of the effect of starvation on growth (Fig 4D). This result corroborates the effect of the double mutant after 8 days of starvation (Fig 4B), except after only 4 days of starvation in this case (4 days of starvation was used since this experiment included a *daf-16* null mutant which is sensitive to starvation). Consistent with our hypothesis, we found no significant difference in the effect of starvation on growth between the null mutant *daf-16(mu86)* and *daf-16(mu86); irld-39(duk1); irld-52(duk17)*, indicating that increased starvation resistance of *irld-39(duk1); irld-52(duk17)* is dependent on *daf-16* (Fig 4D). This example of genetic epistasis suggests that DAF-16 activity is increased in the *irld-39(duk1); irld-52(duk17)* double mutant. As a transcription factor regulated by phosphorylation-dependent subcellular localization, the degree of DAF-16 nuclear localization is a proxy of its activity. In support of our hypothesis, nuclear localization of endogenous DAF-16 (Aghayeva *et al*. 2020) in intestinal cells was significantly increased at 36 hours of L1 starvation in *irld-39(duk1); irld-52(duk17)* mutants (Fig 4E), suggesting that IRLD-39 and IRLD-52 inhibit DAF-16 activity during starvation in wild-type N2 larvae. Together, genetic epistasis and nuclear localization assays suggest that IRLD-39 and IRLD-52 act through DAF-16/FoxO to affect starvation resistance during L1 arrest.

## DISCUSSION

Our results illustrate the power of MIP-seq as a population selection-and-sequencing approach for analysis of complex traits in whole animals. Increasing population complexity and sequencing depth will allow detection of meaningful phenotypic differences too small or variable to be detected by manual assays, leading to improved understanding of gene-by-environment interactions and the genotype-to-phenotype map. With complex traits being highly polygenic (Boyle *et al*. 2017), it will be critical to leverage the power of sequencing to elucidate their architectures. MIP-seq can be used in any organism with known sequence variants and that can be cultured in sufficiently large numbers with the ability to select on the trait of interest.

We characterized natural variation in starvation resistance in a set of genetically diverse, wild strains of *C. elegans* using MIP-seq and traditional assays. Our results suggest relatively little phenotypic variation of this presumably fitness-proximal trait. Nonetheless, we identified the first natural genetic variants affecting starvation resistance in this species. Moreover, the four genes we implicated are all from the same expanded *irld* gene family, suggesting that expansion (or contraction) of gene families influences natural variation and possibly evolutionary adaptation in this context. In addition, two of the *irld* genes identified are in hyper-divergent regions of the genome, consistent with genes in these regions contributing to environmental responses (Lee *et al*. 2021).

Genetic epistasis analysis showed that the effect of *irld-39* and *irld-52* on starvation resistance depends on *daf-16/FoxO*, and we found that nuclear localization of DAF-16/FoxO is affected by *irld-39* and *irld-52.* These observations suggest that these *irld* genes modify IIS, as originally proposed for the gene family (Dlakic 2002). However, *irld-39* and *irld-52* could affect DAF-16 activity independent of IIS. In addition, these *irld* genes could affect other signaling pathways as well. For example, they also bear weak homology to EGF receptors, and *irld* family members *hpa-1* and *hpa-2* affect healthspan by modifying EGF signaling (Iwasa *et al*. 2010). It is not known whether EGF signaling affects starvation resistance or other aspects of L1 arrest, and future work is needed to address the possible role of *irld* genes affecting EGF signaling in this context.

Given the proposal that IRLD proteins modify IIS, it is intriguing to speculate that they do so by binding ILPs that would otherwise bind DAF-2/InsR, as suggested previously (Dlakic 2002). However, it is imperative to emphasize that this is speculation. DAF-2B is an alternative isoform of DAF-2/InsR that includes the extracellular domain but lacks the tyrosine kinase domain, like the IRLD proteins, and it is also thought to act this way (Martinez *et al*. 2020). This hypothetical mechanism is also analogous to the proposed function of IGF-binding proteins, which affect circulation and receptor binding of IGF proteins (Allard and Duan 2018). These parallels suggest the possibility that natural variation in the IGF-binding protein family (Rotwein 2017) contributes to phenotypic variation in humans. The proposed decoy-receptor mechanism could be generalized to other receptor homologs or isoforms to affect a variety of signaling pathways, including deeply conserved pathways. However, we have not shown that IRLD proteins actually bind ILPs, and a variety of uncertainties remain regarding their function in the context of a multicellular animal whose developmental physiology is governed by a complex organismal signaling network.

ILPs have complex spatiotemporal expression patterns and distinct loss-of-function phenotypes (Ritter *et al*. 2013; Fernandes de Abreu *et al*. 2014). ILPs can function as agonists or antagonists of DAF-2/InsR (Pierce *et al*. 2001), and relative expression of these functional classes changes over time during starvation (Chen and Baugh 2014). Assuming that IRLD proteins do bind ILPs, specificity of binding is unclear, but specificity would likely alter the function of individual *irld* genes. In particular, IRLD proteins could differ in binding to DAF-2/InsR agonists or antagonists, or they could bind ILPs indiscriminately, with distinct consequences on IIS in each case. Many but not all IRLD proteins have signal sequences suggesting they are secreted. The extent to which they act locally as opposed to moving freely throughout the body cavity to directly exert systemic effects is unclear. To the extent that IRLD proteins do act locally, their anatomical expression patterns should impact how they affect the IIS network, as well as other signaling pathways they may modify, and possibly organismal phenotype.

Expression analysis provides possible clues to how *irld* genes influence starvation resistance. We used genome editing to knock mScarlet into the *irld-39* and *irld-52* loci to generate endogenous reporter genes, but we were unable to detect their expression during L1 arrest. In an effort to increase sensitivity, we used putative promoter sequences for *irld-39* and *irld-52* to generate transcriptional reporter constructs, and we introduced these as high-copy transgenes, but again we were unable to detect their expression. Published whole-animal mRNA-seq analysis of fed and starved L1 larvae (Webster *et al*. 2018) revealed relatively low expression levels of the entire *irld* family (detected at low levels or not at all) (Fig S6). However, about half of the *irld* genes were differentially expressed, and all of them were expressed at higher levels in starved than fed larvae, suggesting a relatively specific role in starvation. We interrogated two existing single-cell RNA-seq datasets, both of which analyzed expression in fed larvae (either L2-stage (Cao *et al*. 2017) or L4-stage (Taylor *et al*. 2021) larvae). One included each of the major tissue types (Cao *et al*. 2017), and it suggests that *irld* genes are most prominently expressed in ciliated sensory neurons, though there is expression in other types of neurons and other tissues (Fig S7). The other study focused on neurons (Taylor *et al*. 2021), and it also suggests that *irld* gene expression is more prominent in sensory neurons than other neuron types (Fig S7). The authors noted that *irld* family expression is more likely than other gene families to be restricted to a specific neuronal cell type (Taylor *et al*. 2021). *irld-39* is expressed in the ASJ sensory neurons, along with distal tip cells and vulval precursors (Fig S7). *irld-52* is expressed in the ADL sensory neurons and also intestinal rectal muscle cells. *C. elegans* sensory neurons are polymodal and influence life-history traits regulated by IIS, including dauer formation, aging, and L1 arrest (Bargmann and Horvitz 1991; Vowels and Thomas 1992; Apfeld and Kenyon 1999). Furthermore, ASJ is known to express the relatively potent ILP DAF-28 in nutrient and sensory-dependent fashion (Li *et al*. 2003; Kaplan *et al*. 2018), and *daf-28* affects starvation survival (Chen and Baugh 2014). *ins-4* is also expressed in ASJ, and it too affects starvation survival (Chen and Baugh 2014). Together these data suggest that *irld* genes, including *irld-39* and *irld-52*, are expressed in sensory neurons and other, seemingly unrelated cell types. If IRLD proteins are translated and function in the vicinity of these sensory neurons, then that would allow them to exert their influence at the interface of the animal and its environment.

## METHODS

### Strains used in this study

In addition to N2, wild isolates CB4854, CB4856, CX11254, CX11264, CX11271, CX11276, CX11285, CX11307, DL200, DL226, DL238, ED3049, ECA189, ECA191, ECA36, ECA363, ECA369, ECA372, ECA396, ED3017, ED3052, ED3077, EG4724, EG4725, GXW1, JU1212, JU1400, JU1581, JU1652, JU1793, JU1896, JU2001, JU2007, JU2017, JU2106, JU2234, JU2316, JU2464, JU2519, JU2526, JU2576, JU258, JU2592, JU2619, JU2811, JU2829, JU2593, JU2838, JU2841, JU2878, JU2879, JU3137, JU561, JU774, JU775, JU782, KR314, LKC34, MY10, MY16, MY18, MY2147, MY23, MY2453, MY2741, NIC195, NIC199, NIC251, NIC252, NIC256, NIC527, NIC258, NIC261, NIC262, NIC265, NIC266, NIC268, NIC271, NIC3, NIC501, NIC523, NIC526, NIC528, PB306, PS2025, QG2075, QG556, QW947, QX1211, QX1212, QX1791, QX1792, QX1793, QX1794, WN2001, XZ1513, XZ1514, XZ1515, and XZ1516 were phenotyped for starvation resistance using MIP-seq. In addition to these 100 strains, CX11262, ECA348, and NIC260 were included in the MIP-seq pilot but excluded from subsequent analysis based on quality-control metrics described in the ‘MIP-seq analysis’ section. QX1430 was used for validation assays. CB1370 *daf-2(e1370) III,* CF1038 *daf-16(mu86) I,* and OH16024 *daf-16(ot971[daf-16::GFP]) I* were used to assess the interaction of *irld* genes with insulin signaling.

### Strains generated in this study

Near-isogenic lines include:

LRB392 – *dukIR7(V, EG4725>MY2147)*
LRB393 – *dukIR8(V, EG4725>MY2147)*
LRB395 – *dukIR10(V, EG4725>MY2147)*
LRB396 – *dukIR11(V, MY2147>EG4725)*
LRB397 – *dukIR12(V, MY2147>EG4725)*
LRB398 – *dukIR13(V, MY2147>EG4725)*
LRB399 – *dukIR14(V, MY2147>EG4725)*
LRB400 – *dukIR15(V, MY2147>EG4725)*
LRB401 – *dukIR16(V, MY2147>EG4725)*
LRB402 – *dukIR17(V, EG4725>MY2147)*
LRB403 – *dukIR18(V, EG4725>MY2147)*
LRB407 – *dukIR19(V, EG4725>MY2147)*
LRB408 – *dukIR20(V, EG4725>MY2147)*
LRB409 – *dukIR21(V, EG4725>MY2147)*
LRB410 – *dukIR22(IV, DL238>N2)*
LRB411 – *dukIR23(IV, DL238>N2)*

See Figure S3 for wild isolate composition.

CRISPR-edited strains and new crosses include:

LRB412 *irld-39* in N2 background – *irld-39(duk1) IV*
LRB413: *irld-39* in MY2147 background – *irld-39(duk2*[MY2147]) *IV*
LRB414: *irld-39* in MY2147 background – *irld-39(duk3*[MY2147]) *IV*
LRB415: *irld-39* in MY2147 background – *irld-39(duk4*[MY2147]) *IV*
LRB420: *irld-11* in MY2147 background – *irld-11(duk9*[MY2147]) *V*
LRB421: *irld-52* in N2 background – *irld-52(duk10) V*
LRB422: *irld-52* in MY2147 background - *irld-52(duk11*[MY2147]) *V*
LRB423: *irld-11* in N2 background – *irld-11(duk12) V*
LRB425: *irld-57* in N2 background – *irld-57(duk13) V*
LRB426: *irld-57* in N2 background – *irld-57(duk14) V*
LRB427: *irld-57* in MY2147 background – *irld-57(duk15*[MY2147]) *V*
LRB428: *irld-57* in MY2147 background – *irld-57(duk16*[MY2147]) *V*
LRB431: *irld-52* in N2 background – *irld-52(duk17) V*
LRB444: *irld-39; irld-52* (generated from crossing LRB412 and LRB431) - *irld-39(duk1) IV; irld-52(duk17) V*
LRB456: *daf-16(mu86) I; irld-39(duk1) IV; irld-52(duk17) V*
LRB457, LRB458: *daf-2(e1370) III; irld-39(duk1 IV); irld-52(duk17) V*
LRB463: *daf-16(ot971) I; irld-39(duk1) IV; irld-52(duk17) V*

Multiple strain names for the same genotype indicates independent lines.

### MIP-seq experimental set-up

Wild strains were independently passaged on 10 cm NGM plates with OP50 *E. coli* every two to three days to ensure they did not starve for at least three generations prior to the experiment. For each biological replicate, a single non-starved plate with gravid adults was selected per strain to ensure initial representation of all strains. Strains were pooled for hypochlorite treatment to obtain pure populations of embryos (Hibshman *et al*. 2021). Embryo concentration was calculated by repeated sampling, and 500,000 embryos were resuspended at 10/µL in S-basal (50 mL total culture) and placed in a 20°C shaker at 180 rpm to hatch without food and enter L1 arrest. On day 1 (24 hours after hypochlorite treatment), 5 mL of culture (50,000 L1s) was taken as a baseline sample, spun down at 3000 rpm, aspirated down to approximately 100 µL in an Eppendorf tube, flash frozen in liquid nitrogen, and stored at −80°C until DNA isolation. At days 1, 9, 13, and 17, aliquots from the L1 arrest culture were set up in recovery cultures at 5 L1s/µL, 1x HB101 (25 mg/mL), and S-complete. Recovery cultures were 10 mL for days 1 and 9, 20 mL for day 13, and 50 mL for day 17 to account for lethality late in starvation by ensuring adequate population sizes. Four days after recovery culture set-up, samples were collected for DNA isolation. For days 1 and 9, the recovery culture was freshly starved with adults and next-generation L1 larvae. At day 13, the culture was typically near starved, with adults and some L1 larvae. At day 17, the culture was typically not starved. If HB101 was still present at collection, samples were washed 3-4 times with S-basal. Samples were flash frozen in liquid nitrogen and stored at −80°C until DNA isolation.

### DNA isolation

Frozen samples were rapidly freeze-thawed three times, cycling between liquid nitrogen and a 45°C water bath. Genomic DNA was isolated using the Quick-DNA Miniprep Kit (Zymo Research# D3024) following the manufacturer’s protocol. The DNA concentration was determined for each sample using the Qubit dsDNA HS Assay kit (Invitrogen# Q32854).

### MIP design

MIPgen (Boyle *et al*. 2014) was used to design four MIPs for each of 103 strains. Unique homozygous SNVs were parsed from the VCF file WI.20170531.vcf.gz (available at https://storage.googleapis.com/elegansvariation.org/releases/20170531/WI.20170531.vcf.gz). Target regions in BED format were generated using the makeBedForMipgen.pl script. MIPgen was used against *C. elegans* genome version WS245 with the following parameters: -min_capture_size 100 - max_capture_size 100 -tag_sizes 0, 10. MIPs are 80 base-pairs (bp) long and include 20 bp ligation and extension arms that are complementary to DNA surrounding the unique SNV of interest for each strain. In addition, P5 and P7 Illumina sequences are included as part of the MIP to facilitate Illumina sequencing. Each MIP molecule includes a 10 bp unique molecular identifier (UMI) adjacent to the ligation arm. Only MIPs that capture the SNV within a 50 bp sequencing read were used, meaning the SNV was no more than 40 bp away from the UMI. SNPs located within 40 bases of the sequencing start site were parsed with the parseMipsPerSNPposition.pl script. These scripts can be found at https://github.com/amykwebster/MIPseq_2021.

### MIP-seq library preparation and sequencing

For pilots and the starvation-resistance experiment, 500 ng genomic DNA from each sample was used for MIP-seq libraries. Libraries were generated as described previously (Hiatt *et al*. 2013) with the following modifications. We included 1000 copies of each MIP for every individual copy of the worm genome in the 500 ng input DNA, which corresponded to 0.0083 picomoles of each individual MIP. All 412 MIPs (sequences available in Data S1) were first pooled in an equimolar ratio at a concentration of 100 μM. The MIP pool was diluted to 5 μM in 1 mM Tris buffer, and 50 μL of this pool was used in the 100 μL phosphorylation reaction. Next, the probe hybridization reaction for each sample was set with 500 ng DNA and 3.42 picomoles (0.0083 picomoles x 412) of the phosphorylated probe mixture. Following hybridization, gap filling, ligation, and exonuclease steps were performed as described previously. PCR amplification of the captured DNA (primer sequences available in Data S1) was performed in a 50 μL reaction with 18 cycles. The PCR libraries were purified using the SPRIselect beads (Beckman# B23318), and library concentrations were assessed with the Qubit dsDNA HS Assay kit (Invitrogen# Q32854). Sequencing was performed on the Illumina HiSeq 4000 to obtain 50-bp single-end reads.

### MIP-seq analysis

FASTQ files from sequencing reads were processed using the script parseMIPGenotypeUMI.pl, also available at github.com/amykwebster/MIPseq_2021. This script accepts as input the list of MIPs produced from MIPgen, the UMI length, and FASTQ files in order to count the number of reads corresponding to each MIP and whether they have the reference allele, alternative allele, or one of two other alleles. While we included UMIs in our MIP design, use of the UMI to filter duplicate reads in pilot standard curves did not improve data quality (likely due to the relatively large mass of DNA used to prepare libraries), and so the UMI was not used in the published analysis. For each MIP, the frequency of the strain for which the MIP captures its unique SNV was calculated as the alternative read count divided by the total of alternative and reference read counts. For the MIP pilot with all strains in an equimolar ratio, there were 246,986,236 total mapped reads to all MIPs. Individual MIPs were filtered out if they did not meet the following criteria: 1) They were within 3.5-fold of expected frequency (that is, alt / (alt + ref) was within 3.5-fold of 1/103), 2) ‘other’ reads (those that are not alternative or reference alleles) were < 20,000 total, and 3) alternative and reference allele totals were between 20,000 and 2,000,000 total reads. 321 of 412 MIPs met these criteria, and reads from these 321 MIPs were included in subsequent analysis. N2 has very few unique SNVs making it difficult to design optimal MIPs, and N2 MIPs did not meet these criteria but were included nonetheless (see Data S1). For the standard curve experiment (Fig 1F), DNA from seven strains (CB4856, DL200, ED3077, JU258, JU561, JU1652, and N2) was pooled in defined concentrations (‘expected’), and MIPs that met the criteria defined above were used to calculate strain frequencies (‘observed’, see Data S1).

For the starvation-resistance experiment, an average of 51.7 million reads (standard deviation 7.2 million reads) were sequenced per library (one library per time point, replicate, and condition – 25 libraries total). An average of 94% of reads (standard deviation 0.5%) matched the ligation probe, and 71.8% (standard deviation 3.4%) matched the ligation probe and scan sequence. Strain frequencies were determined by averaging the frequencies calculated across MIPs included in the 321 MIPs for each strain. A dataframe of all strains and their frequencies at day 1 baseline, as well as days 1, 9, 13, and 17 after recovery for all replicates was used to obtain trait values for subsequent analysis. PCA was performed on the dataframe following normalization of day 1, 9, 13, and 17 time points by the baseline day 1 sample and log2 transformation. PC1 loadings were extracted for each strain. For ‘Slope’, day 1, 9, and 13 recovery samples were normalized by day 1 frequencies and log2 transformed. For each strain, a line was fit to day 1, 9, and 13 normalized data with intercept at 0, and the slope of the line was taken as the trait value.

### Comparison of MIP-seq and RAD-seq

To determine how well MIP-seq trait values correlated with RAD-seq trait values from previous work (Webster *et al*. 2019), RAD-seq data were normalized the same way that we normalized the MIP-seq data. Specifically, data from one biological replicate from RAD-seq that had data at time points over the course of starvation, including days 1, 7, 14, 21, and 24, was used. The frequency of each strain at each time point was divided by its frequency on day 1. These values were log2 transformed, so positive values indicate an increase in frequency over time and negative values indicate a decrease in frequency over time. A linear regression was then fit to each with a y-intercept of 0 through the data points over time. The slope of the line was calculated as the trait value for each strain. RAD-seq and MIP-seq data were filtered to include only the 34 strains that were present in both analyses. The values were plotted against each other and a linear regression was fit through these points to determine their correlation (R^2^ = 0.24, p = 0.002).

### GWA analysis

Slope and PC1 trait values for each strain were used for GWA using the R package cegwas2 (https://github.com/AndersenLab/cegwas2-nf). Genotype data were acquired from the latest VCF release (release 20200815) from CeNDR. BCFtools (Li 2011) was used to filter variants below a 5% minor allele frequency and variants with missing genotypes and used PINKv1.9 (Purcell *et al*. 2007; Chang *et al*. 2015) to prune genotypes using linkage disequilibrium. The additive kinship matrix was generated from 45,733 markers using the A.mat function in the rrBLUP package. Because these markers have high LD, eigen decomposition of the correlation matrix of the genotype matrix was performed to identify 570 independent tests. GWA was performed using the GWAS function of the rrBLUP package (Endelman 2011). Significance was determined by an eigenvalue threshold by the number of independent tests in the genotype matrix. Confidence intervals were defined as +/- 150 SNVs from the rightmost and leftmost markers passing the significance threshold.

ALT and REF information for *irld-39* and *irld-52* high-impact variants was obtained from fine mapping and is available as part of Data S2. To determine whether *irld-57* and *irld-11* overlapped with hyper-divergent regions in each strain, coordinates of hyper-divergent regions for each strain were obtained from (Lee *et al*. 2021), and coordinates of *irld-11* and *irld-57* were obtained from WormBase. If the hyper-divergent region and gene overlapped for a strain, then the strain was considered hyper-divergent at the locus. Hyper-divergent status of each strain is available in Data S2.

### Enrichment analyses

To identify enriched gene groups, fine mapping data from Slope and PC1 results were merged with WS273 gene names. Unique sequence names were extracted (see Data S2), and the 867 sequence names with variants in Slope or PC1 QTL were used in WormCat (Holdorf *et al*. 2020) to identify functional category enrichments (Fig 3C). The most specific enrichments (those in Category 3) are shown. For genes with unique symbol names (555 genes), those with the same symbol prefix were considered part of the same gene family, and enrichments were calculated for families with at least five members with variants in the QTL. *hpa* and *irld* gene prefixes were grouped under the *irld* prefix (Iwasa *et al*. 2010). P-values were calculated from a hypergeometric test.

For protein domain enrichment analysis, a protein fasta file was downloaded from Wormbase (c_elegans.PRJNA13758.WS281.protein.fa). To determine enriched protein domains, the 867 sequence names present among genes with variants in significant QTL were first used to subset this fasta file. In cases in which a gene had multiple versions within the fasta file, the ‘a’ isoform of the gene was used. 644 of the 867 sequence names had protein sequences in the fasta file; most others are annotated as pseudogenes and presumably do not have protein sequences. The protein sequences were used as input using the hmmscan program (https://www.ebi.ac.uk/Tools/hmmer/search/hmmscan) and searching the Pfam database (Finn *et al*. 2011). To obtain a background set of protein domains, the genome fasta file was also used to search the Pfam database. Fasta files were split into groups of 500 sequences that are between 10 and 5000 peptides to comply with the hmmscan search algorithm. To calculate enrichment of protein domains, hypergeometric p-values were calculated for each protein domain present among genes with variants in significant QTL. 102 protein domains were present, so a Bonferroni-corrected p-value of 0.00049 was used as a significance threshold. Protein domains were excluded if the domain was not present at least five times among genes with variants in significant QTL.

### NIL generation

To validate chromosome IV and V QTL, pairs of strains that differ for starvation resistance and the alternative vs reference allele for the associated SNV marker were chosen. Compatibility at the *peel-1/zeel-1* and *pha-1/sup-35* loci was considered (Seidel *et al*. 2008; Ben-David *et al*. 2017). For chromosome V QTL, EG4725 and MY2147 were compatible at both loci, and we generated reciprocal NILs for the left and right arms of chromosome V. EG4725 did not have the alternative allele associated with starvation resistance for chromosome IV, so we used DL238 and N2 as the parental strains. DL238 and N2 are incompatible for reciprocal crosses, but we introgressed the DL238 chromosome IV QTL into the N2 background. To generate NILs, the two parental strains were first crossed, then F2 progeny were genotyped on each end of the desired QTL for introgression to identify homozygotes from one parental background (*e.g.*, MY2147). Then these homozygotes were repeatedly backcrossed to the opposite background (*e.g.*, EG4725) and repeatedly genotyped to maintain homozygotes at the introgressed region. Genotyping was performed using PCR to amplify a genomic region whose sensitivity to a particular restriction enzyme depends on parental genetic background. Primers were designed using VCF-kit (Cook and Andersen 2017). Primers and enzymes used can be found in Data S2. NILs were backcrossed a minimum of six times. Final NILs were sequenced at ∼1x coverage to determine the parental contributions over the entire genome (Fig S5 and Data S2).

### CRISPR design and implementation to edit irld genes

For genes of interest, CRISPR guide design was done in Benchling using genome version WBcel235 and importing sequence for genes of interest. To generate *irld-39* and *irld-52* variants (5 bp deletions), a single guide RNA (sgRNA) and repair template was generated. For *irld-11* and *irld-57,* two sgRNAs were generated per gene to delete the entire gene and a single repair template was used. sgRNAs (2 nmol) and 100 bp repair templates (highest purity at 4 nmol) were ordered from IDT. The *dpy-10* co-CRISPR method was used to generate and screen for edits (Paix *et al*. 2015). The injection mix used was: sgRNA for *dpy-10* (0.2 µL of 100 µM stock), sgRNA of gene of interest (0.5 µL of 100 µM stock), *dpy-10* repair template (0.5 µL of 10 µM stock), repair template for gene of interest (0.6 µL of 100 µM stock), Cas9 (0.8 µL of 61 µM stock), and water up to 10 µL total. Injection mix components were stored at −20°C, and injection mix was incubated at room temperature for one hour before injections. N2 and MY2147 L4s were picked the day before injecting, and young adults were injected in the gonad and singled to new plates. After 3-4 days, next-generation adults were screened for rollers, which are heterozygous for the *dpy-10* edit and have increased likelihood of also having the desired edit. Non-roller F2 progeny of F1 roller worms were then genotyped to identify worms homozygous for the desired edit, and edits were confirmed by Sanger sequencing. Sequences of sgRNAs, repair templates, and PCR primers for genotyping are available in Data S2.

### Starvation recovery (worm length measurements)

Strains were maintained well-fed for at least three generations prior to beginning experiments. Gravid adults were hypochlorite treated to obtain embryos, which were resuspended at 1 embryo/µL in 5 mL of S-basal in a glass test tube and placed on a roller drum at 20°C so they hatch and enter L1 arrest. After the number of days of L1 arrest indicated on each graph, an aliquot of 500-1000 µL (a consistent volume was used between conditions within the same experiment) per strain was plated on a 10 cm plate with OP50 and allowed to recover for 48 hr. After 48 hr, worms were washed onto an unseeded NGM plate. Images were then taken of worms using a ZeissDiscovery V20 stereomicroscope. To determine lengths of worms, the WormSizer plugin for Fiji was used and worms were manually passed or failed (Moore *et al*. 2013). To determine differences in starvation recovery between strains, a linear mixed-effect model was fit to the length data for all individual worms with duration of starvation and strain as fixed effects and biological replicate as a random effect using the package nlme in R. The summary function was used to calculate a p-value from the t-value.

### Starvation survival

L1 arrest cultures were set up as described for starvation recovery. Starting on the first day of arrest and proceeding every other day, a 100 µL aliquot of culture was pipetted onto a 5 cm NGM plate with a spot of OP50 at the center. The aliquot was placed around the periphery of the lawn, and the number of worms plated was counted. Two days later, the number of worms that had made it to the bacterial lawn and were alive was counted. Live worms that have developed and were outside the lawn were also counted. The total number of live worms after two days was divided by the total plated to determine the proportion alive. For each replicate, a logistic curve was fit to the data, and the half-life (time at 50% survival) was calculated, and a t-test was performed on half-lives between strains of interest. Power analysis was performed in R using the pwr.t.test function in the “pwr” package, with parameters n = 5, sig.level = 0.05, power = 0.5, and type = two.sample. The value of d was calculated, and this was multiplied by the standard deviation of control median survival for that experiment to determine the detectable effect size.

### Early fecundity following starvation

For the assay in Figure 2G, L1 arrest cultures were set up as described for starvation recovery. For the assay in Figure 4C, the experiment was done in a different lab and worms were arrested in a 15 mL conical tube instead of a glass test tube. Conical tubes were rotated continuously at 20°C. For both figure panels, ∼500 L1 larvae were plated on 10 cm OP50 plates at the indicated time point, then allowed to recover for 48 hours. Worms were singled to new 5 cm OP50 plates at approximately 48 hours, then allowed lay progeny until 72 hours, at which time the singled worm was removed. Progeny were counted on these plates 2-3 days later to determine the early fecundity of the individual worms.

### Nuclear localization

L1 arrest cultures were set up as described for starvation recovery. At 36 hours of L1 arrest, an aliquot of 700 µL was spun down in a 1.7 mL Eppendorf tube at 3000 rpm for 30 seconds to pellet L1 larvae. 1.5 µL of worm pellet was pipetted into the center of a slide with a 4% Noble agar pad, and a glass cover slip was immediately placed on top. A timer was set for 3 minutes, and the slide was systematically scanned with each individual worm scored for nuclear localization at 40x or 100x with a Zeiss compound microscope. Nuclear localization of DAF-16::GFP was scored in intestinal cells and assigned as one of four categories: nuclear, more nuclear, more cytoplasmic, and cytoplasmic. ‘More nuclear’ and ‘more cytoplasmic’ are intermediate categories between nuclear and cytoplasmic, with localization closer to being nuclear or cytoplasmic, respectively. Scoring for each slide stopped after 3 minutes. For statistical analysis, nuclear and more nuclear categories were pooled as ‘nuclear’, while cytoplasmic and more cytoplasmic were pooled as ‘cytoplasmic’. The Cochran-Mantel-Haenszel test was used to determine differences in the distribution of the two categories while controlling for biological replicate. See Fig 4E for representative images. DAF-16 is initially very nuclear during L1 starvation, and it moves back to the cytoplasm over time during starvation (Mata-Cabana *et al*. 2020). The 36-hour time point was chosen since it is intermediate in this dynamic process.

### Analysis of published RNA-seq data

We analyzed data from three existing publications (Fig S6, S7) (Cao *et al*. 2017; Webster *et al*. 2018; Taylor *et al*. 2021). First, we re-analyzed whole worm bulk mRNA-seq data from fed and starved N2 L1 larvae (four replicates of each condition from a single batch) (Webster *et al*. 2018). Count data was analyzed using edgeR. 60 *irld* genes were part of the protein-coding gene dataset for genome version WS273, and no minimum expression filter was used to restrict the gene set. The calcNormFactors, estimateCommonDisp, and estimateTagwiseDisp functions were used prior to running the exactTest. An FDR cutoff of 0.05 was used to determine significance. For single-cell data across all major worm tissues (Cao *et al*. 2017),Table S4 from the paper was subset to include only *irld* genes, 63 of which were present in the table. Gene expression is represented in Fig S7 when expression levels of transcripts-per-million are at least 1 for that tissue type. For single-cell neuronal data (Taylor *et al*. 2021), expression values for *irld* genes were obtained from Supplementary Table 11 of the published paper, which includes genes considered expressed at a variety of thresholds and neuronal cell types. Data plotted in Figure S7 uses threshold 3 and data from sensory neurons.

### Data availability

Raw MIP-seq data for the starvation-resistance experiment and the pilot experiments to test individual MIPs is available as part of NCBI BioProject PRJNA730178. Code for processing MIP-seq data is available at github.com/amykwebster/MIPseq_2021.

## Acknowledgments

We thank Oliver Hobert for providing OH16024 *daf-16(ot971[daf-16::GFP])*, Jon Hibshman for sharing a starvation survival curve-fitting script, Chelsea Shoben for help passaging wild isolates, Clay Dilks for CRISPR advice, Sophia Gomez for genotyping assistance, Seth Taylor for strain organization and maintenance, and Jim Jordan for helpful discussions. Funding was provided by the NIH (R01GM117408 and R01GM143159 to LRB and R01ES02993 to ECA and LRB). AKW was supported by an NSF Graduate Research Fellowship. Some strains were provided by the CGC, which is funded by NIH Office of Research Infrastructure Programs (P40 OD010440). We would also like to thank WormBase.

## Author contributions

Conceptualization: AKW, LRB

Investigation: AKW, RC, MP, JC, KF, RT, AW

Formal analysis: AKW, LS, KE, IA, AW

Visualization: AKW

Funding acquisition: LRB, ECA

Supervision: LRB, ECA

Writing – original draft: AKW, LRB

Writing – review & editing: AKW, LRB, ECA, RC, KE, IA

## Materials and correspondence

Correspondence and material requests should be addressed to.

## SUPPLEMENTARY FIGURES

**Fig. S1.**
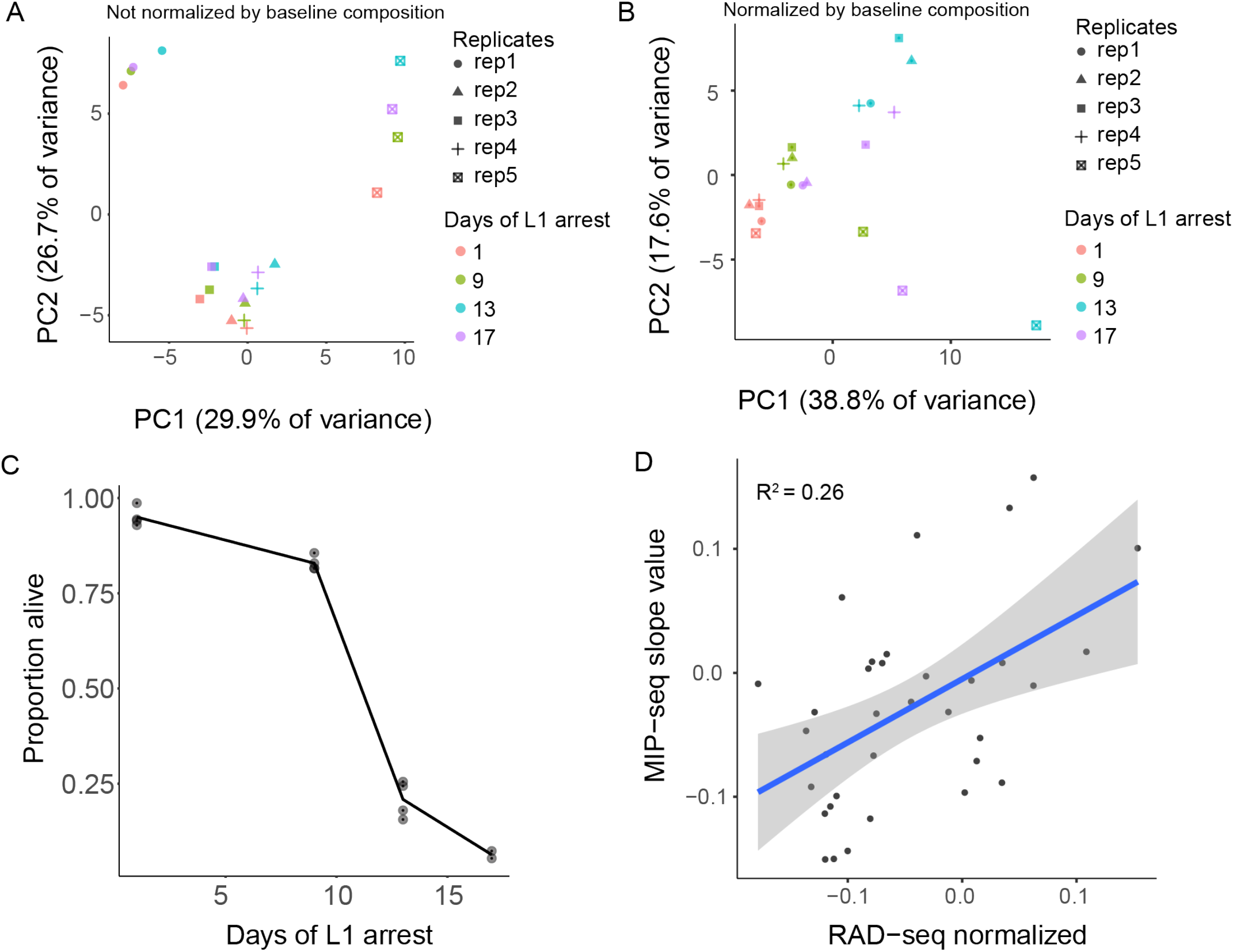
MIP-seq data analysis and comparison to RAD-seq analysis of starvation resistance. (**A**) PCA of all replicates and recovery time points assayed not normalized by baseline composition. Samples cluster by replicate. (**B**) PCA of all replicates and recovery time points normalized by baseline composition of each replicate pool on day 1 of starvation without recovery (See Fig. 2A). PC1 is plotted in Fig. 2B. (**C**). Survival of cultures used to collect samples for MIP-seq experiment (Fig. 2) at days 1, 9, 13, and 17. (**D**) Comparison of MIP-seq trait value Slope to similar trait value derived from published results using RAD-seq for population sequencing (Webster *et al*. 2019). 34 strains included in both experiments are plotted. RAD-seq results are based on only a single replicate, and the vast majority of reads from RAD-seq do not cover unique SNV greatly limiting sequencing depth at informative loci. R^2^ = 0.26 (p = 0.002).

**Fig. S2.**
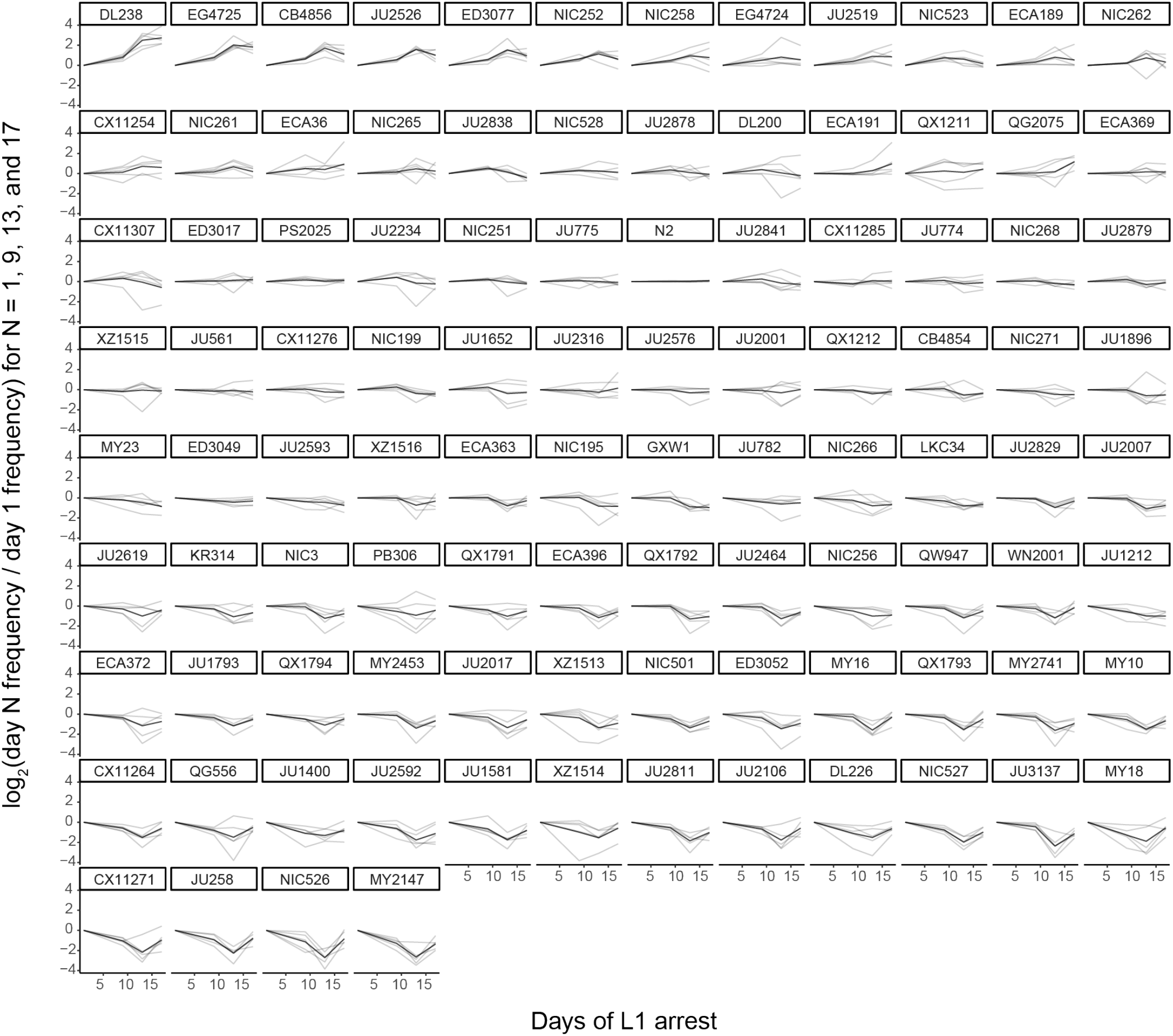
MIP-seq strain frequencies throughout starvation for all strains assayed. MIP-seq replicates were first normalized by baseline and then by day 1. Strains are rank-ordered from most starvation-resistant to most sensitive by Slope trait values. Gray lines are individual replicates and black lines are the mean for each strain.

**Fig. S3.**
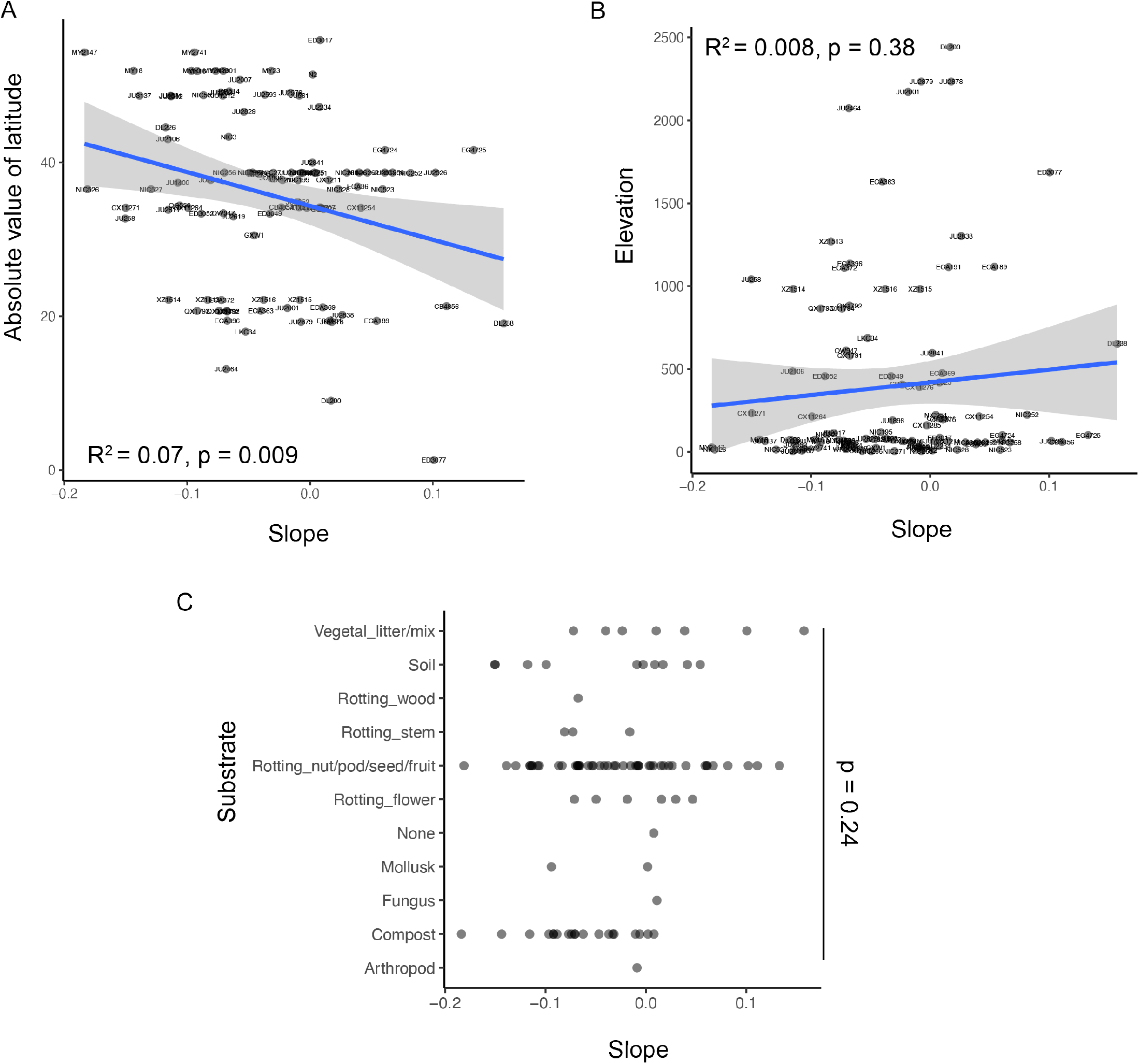
Starvation resistance of wild strains is associated with latitude at collection site, but not elevation or substrate. (**A**) ‘Slope’ trait value for each strain plotted against the absolute value of its latitude of collection. R^2^ = 0.07, p = 0.009. (**B**) ‘Slope’ trait value for each strain plotted against the elevation from which strains were collected. R^2^ = 0.008, p = 0.38. (**A-B**) Significance determined by t-test on t-statistic of slope coefficient from linear model. (**C**) Slope trait values plotted for strains collected from each type of substrate. One-way ANOVA across all substrates, p = 0.24.

**Fig. S4.**
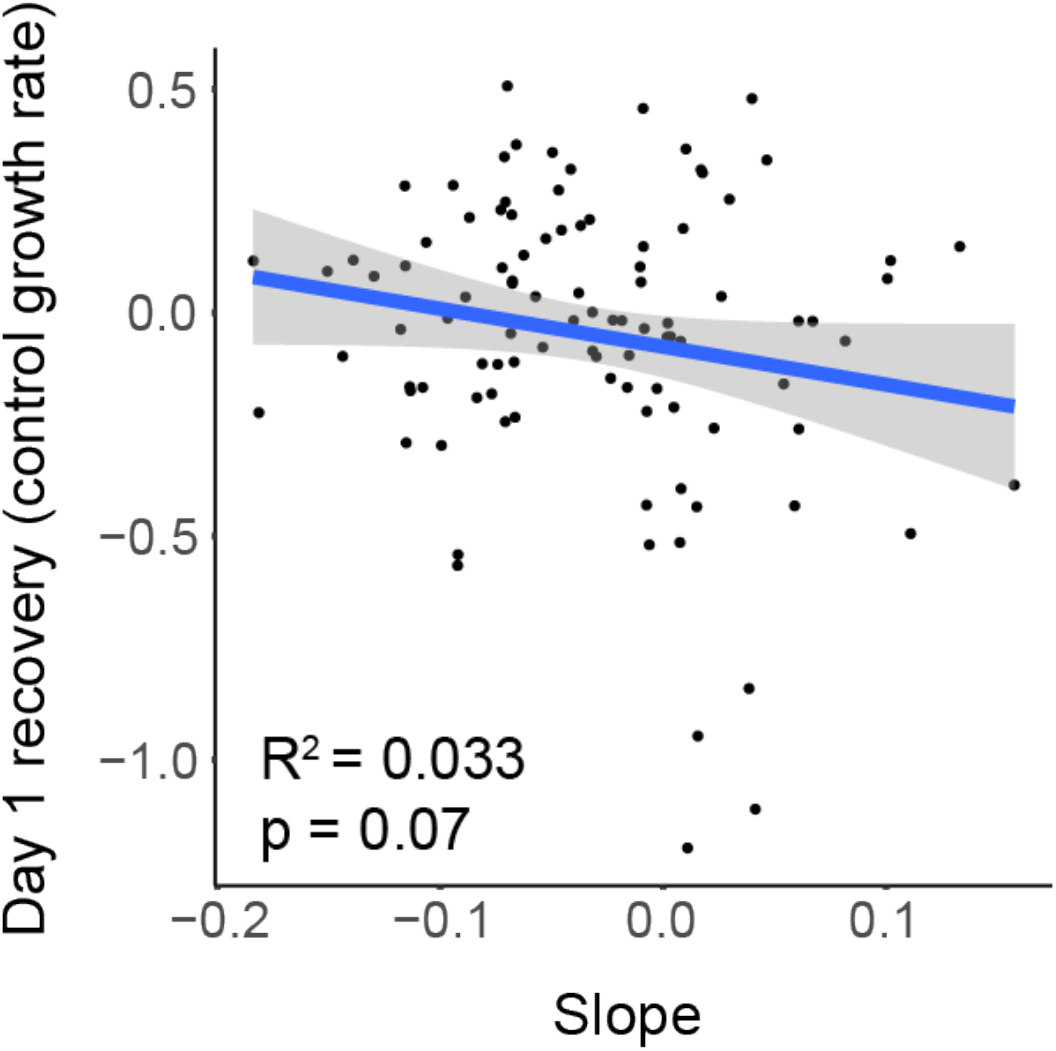
Starvation resistance is negatively correlated with growth rate. Slope trait value is plotted against ‘Early Growth’ trait value calculated by determining which strains became over- or under-represented after recovery from day 1 of L1 starvation relative to baseline composition at day 1 of L1 starvation. R^2^ = 0.033, significance determined by t-test on t-statistic of slope coefficient from linear model.

**Fig. S5.**
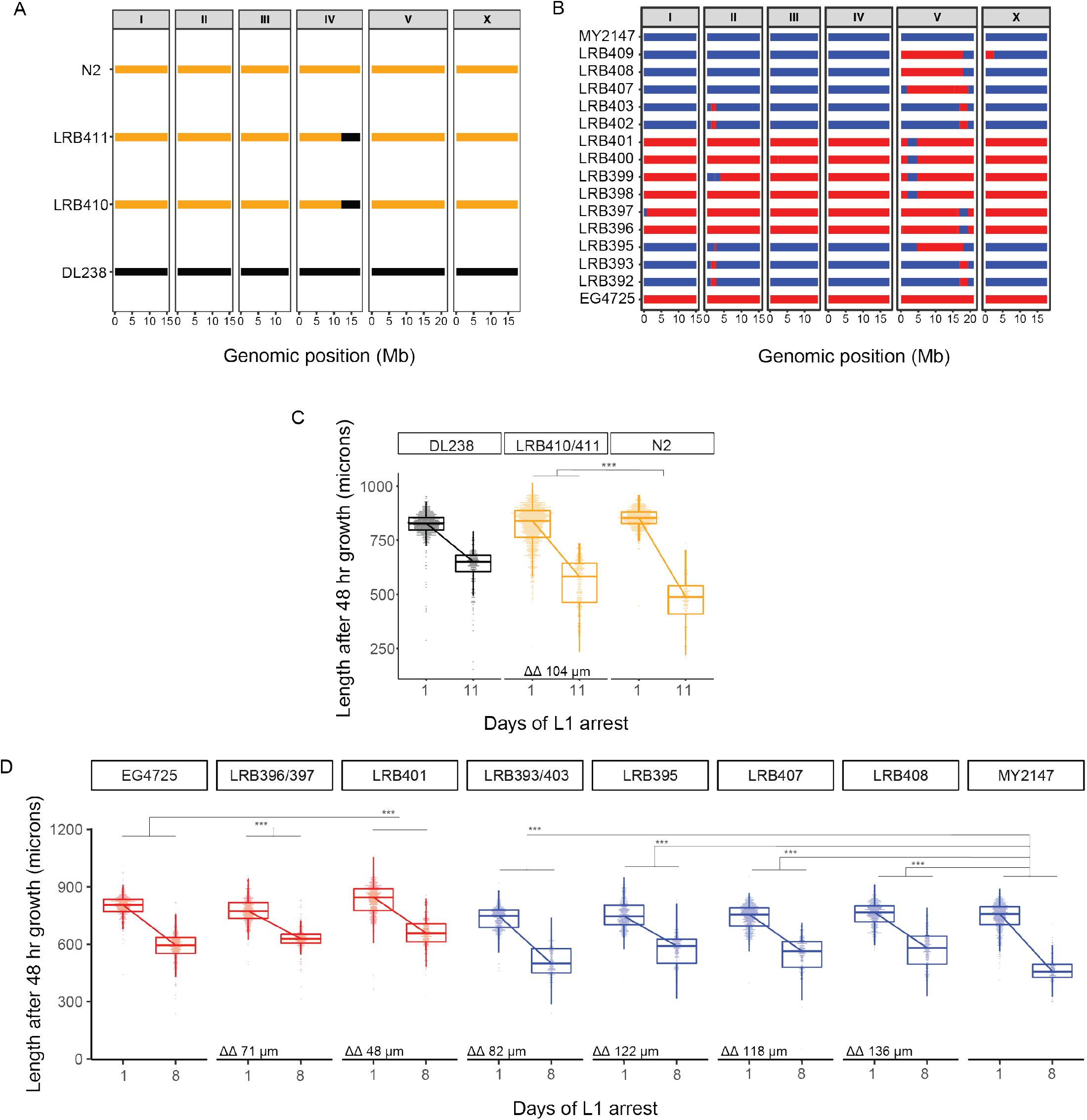
Validation of QTL with sequenced near-isogenic lines (NILs). (**A**) Schematic of low-depth (∼1x) genome-sequencing results of NILs used to validate chromosome IV QTL. Parental strains are N2 and DL238. DL238 can be introgressed into N2 but not *vice versa* due to genetic incompatibility (Ben-David *et al*. 2017). N2 sequence is in gold, while DL238 sequence is in black. (**B**) Schematic of low-depth (∼1x) genome-sequencing results of NILs used to validate chromosome V QTL. Parental strains are MY2147 and EG4725, which are compatible and were introgressed reciprocally. MY2147 sequence is in blue, while EG4725 sequence is in red. (**C**) Worm length following 48 hr recovery with food after 1 or 11 d of L1 starvation for indicated strains. Data were merged for essentially duplicate strains LRB410 and LRB411. Color-coding indicates the primary parental background (*e.g.,* LRB410 is primarily the N2 background so is plotted in yellow). (**D**) Worm length following 48 hr recovery with food after 1 or 8 d of L1 starvation for indicated strains. Data were merged for essentially duplicate strain pairs LRB396 and LRB397, and LRB393 and LRB403. Color-coding indicates the primary parental background (*e.g.,* LRB401 is red because its sequence is primarily EG4725). (**C-D**) ***p < 0.001; linear mixed-effects model; p-value of interaction between strain and duration of starvation; eight biological replicates. ΔΔ indicates effect size of interaction between duration of starvation and strain data plotted in that panel compared to the predominant parental strain background (the difference in differences between strains’ mean length at days 1 and 8).

**Fig. S6.**
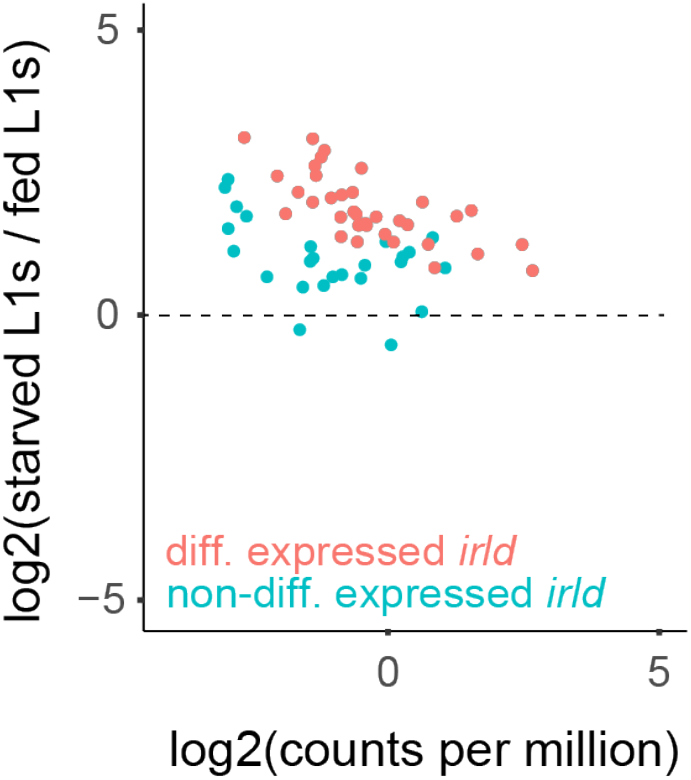
*irld* genes are up-regulated in starved L1s compared to fed L1s. Starved and fed L1 mRNA-seq data from whole, wild-type (N2) worms were re-analyzed from Webster et al. (2018) without a minimum expression filter. *irld* genes are expressed at low levels (typically below one count-per-million reads) but exhibit a consistent pattern of up-regulation in starved compared to fed L1s. Color-coding indicates significance at FDR < 0.05. “diff.” differentially; “non-diff.” non-differentially.

**Fig. S7.**
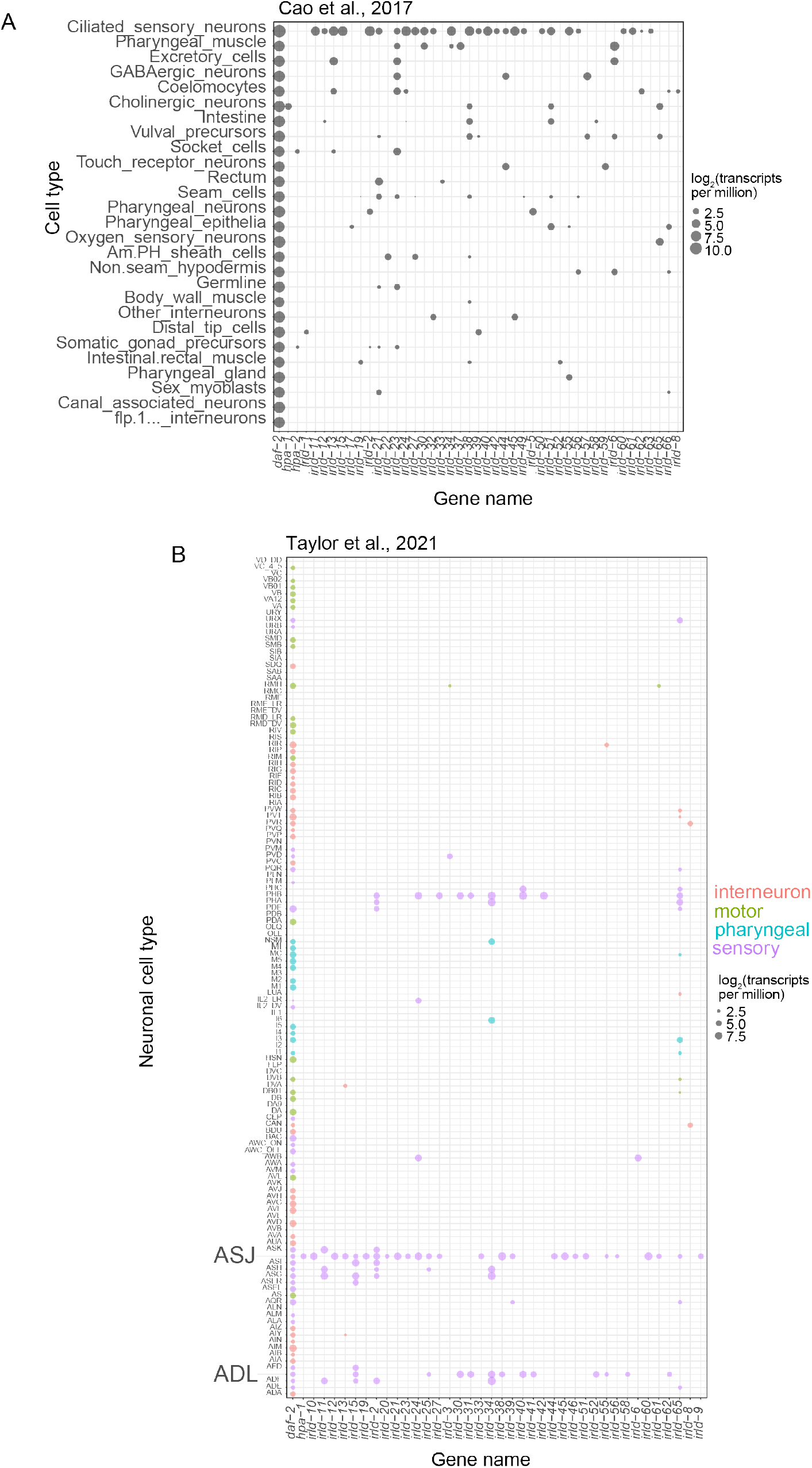
*irld* genes exhibit a bias toward expression in sensory neurons with some expression in other cell types. Single-cell RNA-seq expression levels for all detectable *irld* genes and the insulin/IGF receptor *daf-2* included for comparison. (**A**) Expression levels (transcripts-per-million) from L2-stage larvae for all available cell types are plotted (Cao *et al*. 2017). (**B**) Expression levels from L4 larvae for all neurons (Taylor *et al*. 2021) are plotted. Neurons are sorted into interneuron, motor, pharyngeal, and sensory classifications. Bubble size indicates log_2_ expression level in transcripts-per-million.

**Data S1. (Data_S1.xlsx)**

This file includes all MIP-seq processed data: Slope and PC1 trait values used in GWA, output from MIPgen, MIP sequences, MIPs included in the final experiment, MIP primer sequences, count data for MIP-seq starvation resistance experiment and two pilot experiments.

**Data S2. (Data_S2.xlsx)**

This file includes GWA output and follow-up on *irld* candidates: GWA output for both Slope and PC1, genes within QTL, output from WormCat and protein domain enrichment analyses, hyper-divergence status for each strain, CRISPR sequences, genotyping primers, and NIL sequencing results.

**Data S3. (Data_S3.xlsx)**

This file includes input and output for RNA-seq analysis, including count tables and differential expression output from Webster et al. (2018) used in Fig S6, transcripts-per-million from Cao et al. (2017) plotted in Fig S7, and transcripts-per-million for threshold 3 from Taylor et al. (2021) plotted in Fig S7.

